# Previously uncharacterized aliphatic amino acid positions modulate the apparent catalytic activity of the EAL domain of ZMO_1055 and other cyclic di-GMP specific EAL phosphodiesterases

**DOI:** 10.1101/2024.06.21.600002

**Authors:** Lianying Cao, Xue Zhang, Feng-wu Bai, Ute Römling

## Abstract

The ubiquitous second messenger cyclic di-GMP signaling system decides bacterial life style transition between sessility and motility. GGDEF diguanylate cyclase and EAL phosphodiesterase domains conventionally direct the turnover of this signaling molecule being subject to micro- and macroevolution. While the highly conserved signature amino acids involved in divalent ion binding and catalysis have readily been identified, recognition of amino acid substitutions that modulate the catalytic activity is rare. Associated with development towards cellulose-mediated self-flocculation in *Zymomonas mobilis* ZM401, the A526V substitution, previously not been recognized to affect the functionality of the EAL domain, downregulates the apparent catalytic activity of the PAS-GGDEF-EAL ZMO1055 phosphodiesterase compared to parental *Z. mobilis* ZM4 and equally in homologous protein domains independently of the genetic background. Substitution of A526, which is conserved among homologs, with amino acids with longer aliphatic side chains than valine have an even more pronounced effect. Thus single amino acid substitutions that lead to alterations in the catalytic activity of cyclic di-GMP turnover domains amplify the signaling output and thus significantly contribute to the flexibility and adaptability of the cyclic di-GMP signaling network. In this context, ZMO1055 seems to be a current evolutionary target.

## Introduction

To date, a vast number of cyclic nucleotide-based second messenger molecules have been identified in bacteria (Ross, Weinhouse et al., 1987, Whiteley, Eaglesham et al., 2019, Witte, Hartung et al., 2008). Remarkably, some of them were predicted earlier by docking experiments or had been already chemical synthesized (de Vroom, Broxterman et al., 1988, Hsu &Dennis, 1982, Ross, Mayer et al., 1990). However, among these molecules, cyclic di-GMP seems to be (one of) the most widespread molecule with abundant presence (Romling, 2023, Romling, Galperin et al., 2013). Synthesized by members of the GGDEF domain superfamily and hydrolyzed by members of the EAL domain superfamily, the complexity of the cyclic di-GMP network is reflected by the multiplicity of cyclic di-GMP turnover proteins and structural diversity of receptors harbored by an individual bacterial genome (Romling, Gomelsky et al., 2005, Schirmer &Jenal, 2009). As all domain superfamilies, the GGDEF and EAL superfamilies are subject to macro- and microevolution (Beyhan &Yildiz, 2007, Romling, Cao et al., 2023) with catalytically inactive and/or substrate or product binding receptor domains to be frequently identified. Intriguingly, ongoing microevolution, may it be in the laboratory setting or observed in closely related isolates or species, allows the identification of evolutionary mechanisms and initial alterations in the amino acid sequence of protein evolution in almost real time (Cimdins, Simm et al., 2017, Farr, Remigi et al., 2017, Zlatkov &Uhlin, 2019). For example, originally identified as a catalytically incompetent protein and characterized as a small RNA binding protein in commensal *Escherichia coli*, the GAPES4-HAMP-GGDEF-EAL protein CsrD can exhibit cyclic di-GMP binding and even catalytic activity in other species of the Enterobacteriaceae (Fineran, Williamson et al., 2007, Kharadi &Sundin, 2022, Potts, Leng et al., 2018). Equally, three most closely related EAL only domain proteins encoded by the *Salmonella typhimurium* genome either display catalytic activity or work exclusively through protein-protein interactions (Ahmad, Wigren et al., 2013, El Mouali, Kim et al., 2017).

*Zymomonas mobilis* is an organism with a high potential application in industry for lignocellulosic ethanol production (Xia, Yang et al., 2019). Selection of the *Z. mobilis* ZM4 strain for enhanced ethanol tolerance created the self-flocculating strain *Z. mobilis* ZM401 (Lee, 1982, Zhao, Bai et al., 2014). Sequencing of this strain identified 32 single nucleotide polymorphisms and one single nucleotide deletion in a poly-nucleotide tract (Cao, Yang et al., 2022, Jeon, Xun et al., 2012, Zhao et al., 2014). Thereby, considered most relevant for the self-flocculating phenotype were the single nucleotide deletion that lead to the fusion of the ZMO1082 and ZMO1083 genes resulting in a longer BcsA cellulose synthase open reading frame and gene product. On the other hand, the nonsynonymous single nucleotide polymorphism C1577T created the A526V amino acid substitution in the EAL domain of the GGDEF-EAL domain protein ZMO1055_ZM4_. ZMO1055 A526V, subsequently designated ZMO1055_ZM401_, displayed a lower apparent phosphodiesterase activity and highly reduced capacity to dissolute the self-flocculation phenotype caused by elevated cellulose production in strain *Z. mobilis* ZMO401 (Cao et al., 2022, Jeon et al., 2012). Indeed, deletion and overexpression analysis in combination with mutant construction of all cyclic di-GMP turnover proteins in *Z. mobilis* ZM4 and ZM401 showed that ZMO1055 is a key diguanylate cyclase/phosphodiesterase regulating cellulose triggered self-flocculation (Li, Xia et al., 2023a).

Recognizing the impact of the A526V substitution located in an amino acid stretch of the EAL domain not previously identified as functionally relevant, we were asking whether the impact of the A526V substitution is observed in other model systems and whether the amino acid position equivalent to A526 in ZMO1055 is conserved among the EAL domains and whether functional conservation of the A526V substitution occurs. We found that alanine is predominant at the 526 position equivalent in ZMO1055 homologs and other EAL domain proteins and that the substitution of alanine 526 and upstream position 525 by amino acids with a more bulky aliphatic, polar or aromatic side chain mediates a functionally conserved behavior in more than one subgroup of EAL domain proteins.

## Experimental Procedures

### Strain and growth conditions

The strains used in this work are listed in Table S1. *E. coli* and *S. typhimurium* strains were cultivated in Luria-Bertani (LB: 1% tryptone, 0.5% yeast extract, 1% NaCl) medium, *Z. mobilis* ZM4 strains and derivatives were cultivated in Rich-Medium (RM, 1% yeast extract, 2% glucose, 0.2% KH_2_PO_4_) starting with an initial OD_600_ = 0.1. Strains containing the pEZ plasmid were cultivated in medium supplied with 100 mg/L spectinomycin, strains harboring the pBAD28 or pBAD30 plasmids were supplied with 25 mg/L chloramphenicol and 100 mg/L ampicillin, respectively. 100 ng/L anhydrotetracycline and 0.1% L-arabinose were applied to induce gene expression from the pEZ and pBAD plasmids, respectively.

### Plasmid construction

The pEZ backbone was constructed by Golden Gate cloning (Engler, Kandzia et al., 2008). The plasmid backbone and the P_aTc_ promoter were amplified from plasmid pEZ-dual promoter (Yang, Shen et al., 2019). Gene ZMO1055_ZM4_ or ZMO1055_ZM401_ were amplified from the genomes of *Z. mobilis* ZM4 and ZM401. Plasmid backbone, promoter and gene fragments were purified by Cycle Pure (Omega), restricted with BsaI (NEB Biolabs) and then ligated with T4 DNA ligase (NEB Biolabs) using the Golden Gate cloning protocol (Engler et al., 2008). Recombinant plasmids were transformed into *E. coli* K12 TOP10 and confirmed by PCR and sequencing. Verified plasmids were transformed into *S. typhimurium* UMR1 Δ*yhjH* or *Z. mobilis* ZM401 for assessment of biofilm morphotypes, self-flocculation and motility. Plasmids used and constructed in this work are provided in Table S2. Primers are found in Table S3.

### Site-directed mutagenesis

Site-directed mutagenesis was performed by the Q5 site-directed mutagenesis kit (NEB) with primers designed to harbor the mutation site(s). Plasmids obtained with the help of this kit were confirmed by PCR and sequencing. Verified plasmids were transformed by electronporation (Biorad Gene Pulser^TM^) into their respective hosts for *in vivo* assessment of protein function.

### Congo red binding assay for Z. mobilis

The Congo red binding was pictured to reflect the expression of cellulose in *Z. mobilis*. Shortly, a loopful of bacteria was collected from plate and inoculated into rich medium containing the suitable antibiotic with initial OD_600_ = 0.1. After cultivating at 30 for 12 hours, the bacteria were collected and re-suspended in RM and subsequently adjusted to OD_600_ = 5.0 as seed culture. Three μl of seed culture were inoculated into Congo red plate (2% glucose, 1% yeast extract, 0.2% KH_2_PO_4_, 40 μg/ml Congo red, 20 μg/ml Coomassie Brilliant Blue G-250, 1.5% agar) and incubated at 30 °C. The Congo red morphotype was documented after 24 h and 48 h.

### Calcofluor white colony morphotype assay for Z. mobilis

Calcofluor white (fluorescent brightener 28) binding was visualized to reflect the expression of cellulose. A seed culture was prepared as described for the Congo red assay. Three μl of seed culture was inoculated onto a calcofluor white plate (2% glucose, 1% yeast extract, 0.2% KH_2_PO_4_, 50 μg/ml Fluorescent Brightener 28, 1.5% agar) and cultivated at 30 °C. The calcofluor white binding was documented after 24 h.

### S. typhimurium swimming assay

Swimming motility was quantified by the diameter of the swimming halo (Li, Cimdins et al., 2019). Shortly, a single colony was picked and inoculated onto an LB medium plate containing the suitable antibiotic. After incubation of the plate at 30 ℃ overnight, the bacteria were re-suspended in LB without NaCl medium and subsequently adjusted to OD_600_ = 5.0 as seed culture. Three μl of seed culture was inoculated into soft agar LB medium (1% tryptone, 0.5% yeast extract, 1% NaCl, 0.25% agar) and incubated at 30 °C. The swimming diameter was documented each hour from 5 to 9 hs.

### Rdar colony morphotype assay for S. typhimurium

The rdar morphotype to reflect the expression of cellulose and curli fimbriae was visualized as follows. A seed culture was prepared as described for the swimming motility assay. Three μl of seed culture was inoculated onto rdar plate (1% tryptone, 0.5% yeast extract, 40 μg/ml Congo red, 20 μg/ml Coomassie Brilliant Blue G-250, 1.5% agar) and incubated at 30 °C. The rdar morphotype was documented after 24 h.

### Western blot analysis

To detect protein production, 5 mg of *Z. mobilis* cells were collected from a liquid culture by centrifugation, resuspended in 200 μl SDS sample buffer and heated at 95 °C for 10 min. For *S. typhimurium* strains, 5 mg of cells were collected and re-suspended in 100 μl SDS sample buffer. The protein concentration was assessed by Coomassie Brilliant blue staining after running of the protein extracts on an SDS-PAGE (4% stacking and 12% resolving gel). Samples normalized for the protein content were separated by SDS-PAGE and transferred onto a PVDF membrane (Millipore). After washing with TBS buffer twice, the membrane was blocked in Anti·His HRP Conjugate blocking buffer (Penta·His HRP Conjugate, Qiagen) at 4 °C overnight. The following steps were conducted at the manufacturer’s (Qiagen) instructions with 1:2000 dilution of the antibody. Chemiluminescent detection reaction was done by the ECL light detection reagent (Roche) using Luminescent Image Analyzer (LAS-1000plus, Fujifilm).

### Bioinformatic analyses including protein alignment and construction of the phylogenetic tree

5039 proteins most similar to ZMO1055 as retrieved by BLAST search (April 2021) from the NCBI database and all other relevant GGDEF/EAL proteins were aligned with ZMO1055_ZM4_ by Clustal 2.1 (Higgins &Sharp, 1988) and manually curated in Genedoc. Construction of the phylogenetic relationship and assessment of robustness of the GGDEF and EAL domains was done with Evolview with Bootstrap setting below 50 as grey nodes and for 51-100 as black nodes. All other phylogenetic trees (Maximum-Likelyhood with 1000 bootstraps iterations) were constructed and visualized with MEGA 7.0 or 11.0 (Kumar, Stecher et al., 2016). The DGC activity of GGDEF domains of proteins was discriminated after bioinformatic analysis in two different classes: GGDEF domains with all amino acid signatures conserved with functional reference GGDEF domains were classified as DGC active domain; domains with any amino acid altered in functionally relevant signature motifs were classified as uncertain to possess DGC activity. For PDE activity, EAL domains with all signature amino acids conserved with reference catalytically functional domains were classified as PDE active domains, domains with the catalytic base glutamine in the EGVE motif not conserved were classified as PDE inactive domains and the other domains are classified to be uncertain to possess PDE activity.

Sequence logos of aligned domains have been constructed with WebLogo 2.0 (Crooks, Hon et al., 2004, Schneider &Stephens, 1990). Aligned sequences have been visualized with ESPript 3.0 (Robert &Gouet, 2014). EasyFig (Sullivan, Petty et al., 2011) has been used to visualize alignments of sequence contigs using standard parameters. Structural models for ZMO1055 have been created with Phyre2 (Kelley, Mezulis et al., 2015), Swissprot or I-Tasser (Yang, Yan et al., 2015). The Phyre2 highest scoring templates derived for the ZMO1055 GGDEF domain from PA0575, PA0861 and PA1120 and for the EAL domain from MorA (PA4601), PA0575 and BifA, all from *P. aeruginosa*.

## Results

### Initial characterisation of the GGDEF/EAL domain proteins of Zymomonas mobilis ZM4

The *Zymomonas mobilis* ZM4 genome codes for five GGDEF/EAL domain containing proteins. In order to associate their catalytic capability with sequence conservation, the conserved sequence motifs were analyzed after comparison of the domain alignment with functional DGCs/PDEs. Four proteins, ZMO0919, ZMO1365, ZMO0401 and ZMO1055, contain a GGDEF domain (Figure S1a and 1a; (Li et al., 2023a)). The GGDEF domains of ZMO0919 and ZMO1365, possess all signature amino acid motifs except residue L29 which was substituted by I and M for ZMO0919 and ZMO1365, respectively. Leucine, isoleucine and methionine are all non-polar amino acid and have similar size of side chain. Substantial DGC activity of ZMO0919 and ZMO1365 has been observed as predicted (Li et al., 2023a).

The GGDEF domains of ZMO0401 and ZMO1055, however, possess a degenerated GGDEF motif altered to GNDEF and GGDQF, while the substrate interacting residue D191 and the transition state stabilizing residue K194 are not conserved in ZMO1055_ZM4_ as in the GGDEF domains of highly similar GGDEF-EAL domain proteins (Figure 1a). We have, however, recently shown that the GNDEF/GGDQF domains of ZMO0401 and ZMO1055 still possess diguanylate cyclase catalytic activity (Li et al., 2023a).

**Figure 1:**
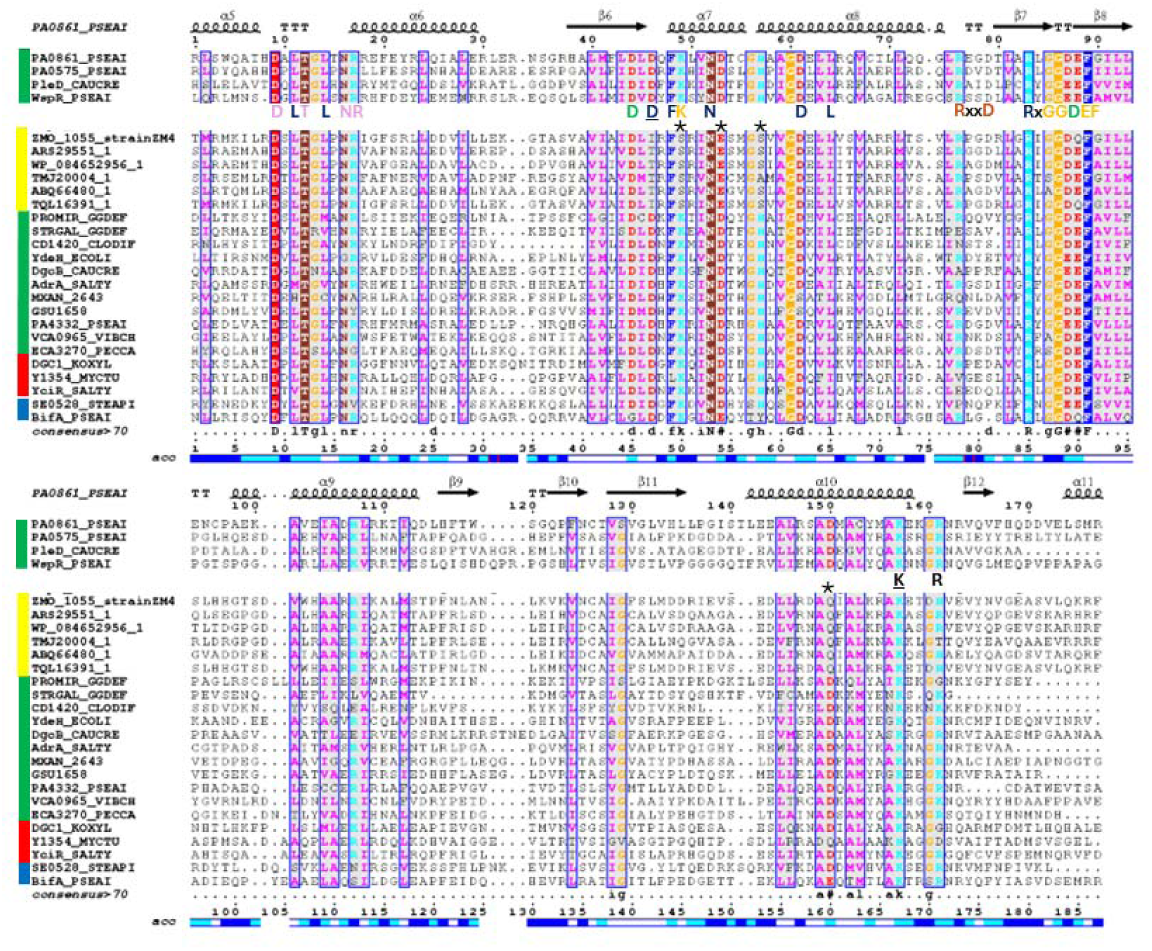
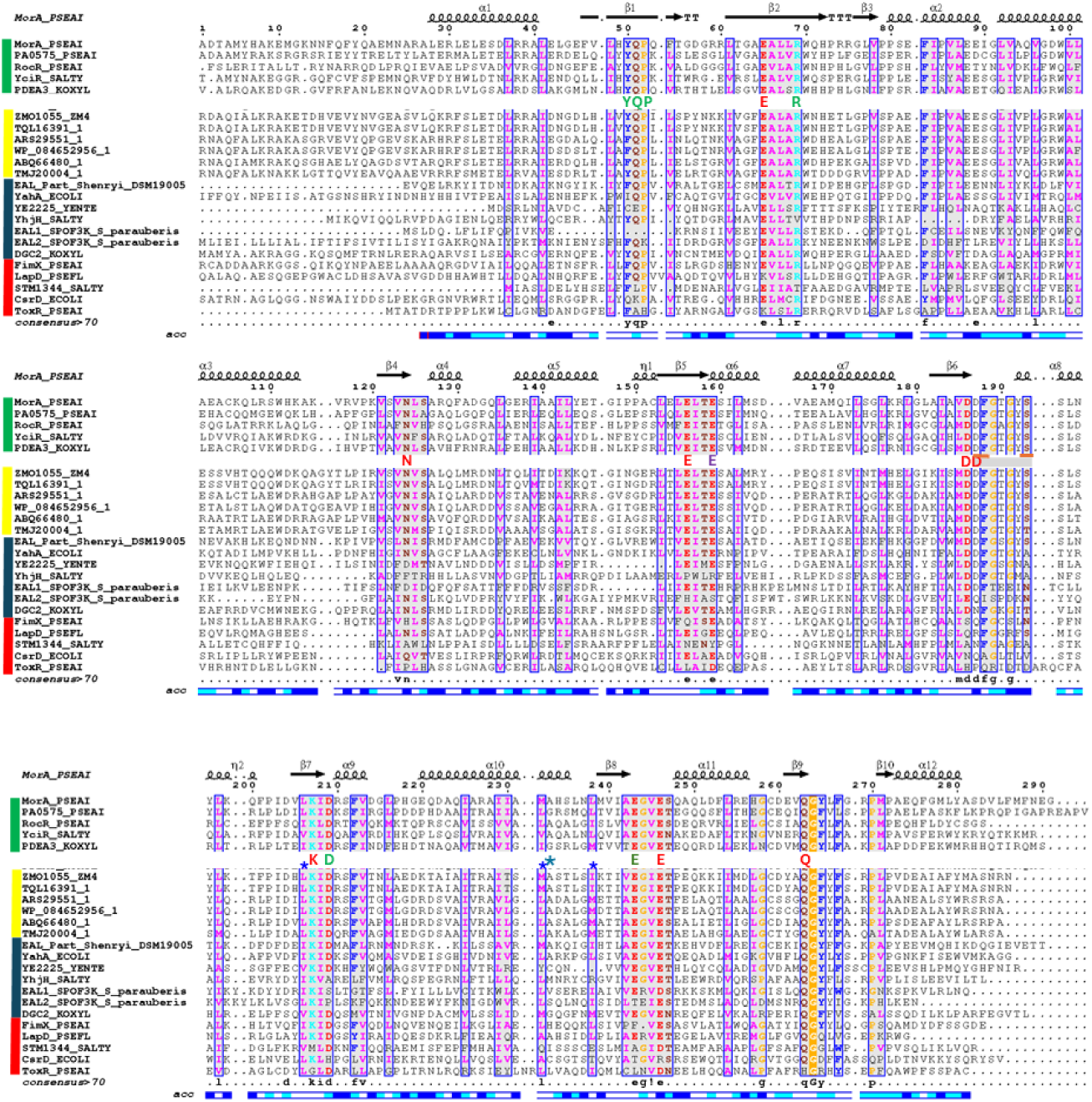
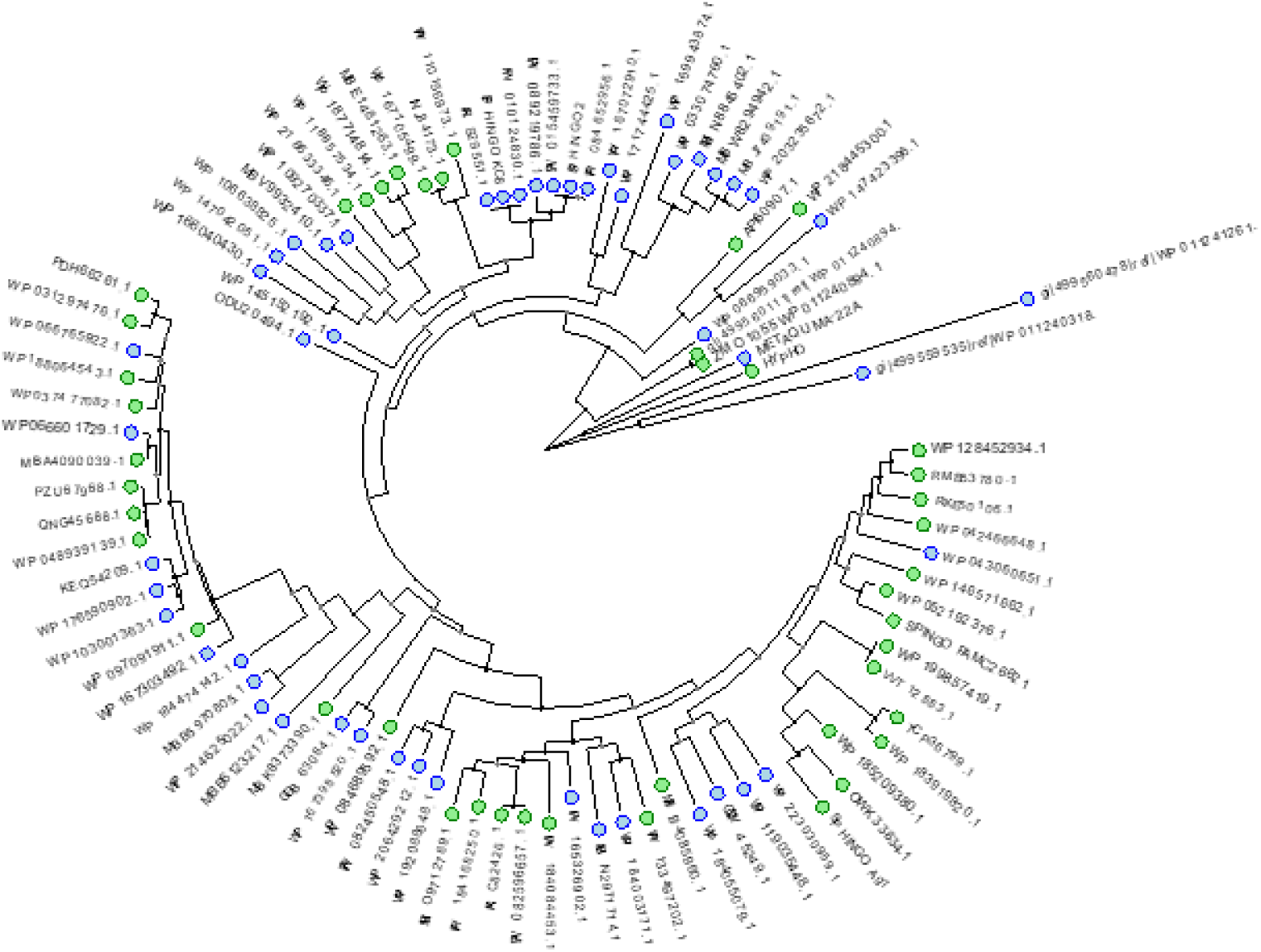
Alignment of the diguanylate cyclase/phosphodiesterase ZMO1055_ZM4_ GGDEF and EAL domains with the respective domains of most similar proteins and reference GGDEF and EAL domains. Figure 1a. Alignment of the ZMO1055_ZM4_ GGDEF domain with functionally characterized GGDEF domains (Liu et al., 2020) and GGDEF domains from most similar GGDEF-EAL domain proteins (see Figure S4e; yellow bar) The GGDEF domain of the GGDEF-EAL protein PA0861 (PDB: 5XGD) has been taken as the highest scoring template as reference for secondary structure-sequence correlation. Functionally assigned amino acids from conserved signature motifs are indicated below the PleD (*Caulobacter vibrioides*) and WspR (*Pseudomonas aeruginosa*) reference amino acid sequences. Stars indicate amino acids not conserved in the GGDEF domain of ZMP1055 and its homologs. Functionality of amino acids in light blue, wide turn in protein; in dark blue, substrate interacting residues; in green, Mg^2+^ binding; in dark yellow, stabilizing the transition state; conserved in plum, allosteric I-site; GG[D/E]EF motif in yellow; underlined, salt bridge. Selected demonstrated catalytically functional class I GGDEF domains (indicated by green bar), catalytically functional class II GGDEF domains of GGDEF-EAL proteins (indicated by red bar) and catalytically non-functional class III GGDEF domains (indicated by blue bar) have been used in the alignment. Figure 1b. Alignment of ZMO1055_ZM4_ EAL domain with functionally characterized EAL domains and EAL domains from most similar GGDEF-EAL domain proteins (see Figure S2e; yellow bar). The EAL domain of the phosphodiesterase MorA (PA4601) (PDB: 4RNJ) as highest scoring template in Phyre2 modelling has been taken as reference for secondary structure-sequence correlation. Green star indicates amino acid A526 and blue stars indicate amino acids 499, 525 and 531 substituted in the course of this work. Functionally assigned amino acids from conserved signature motifs are indicated below the PA0861 and RocR reference amino acid sequences. Functionality of amino acids in green, amino acids involved in substrate binding; in red, amino acids involved in Mg^2+^ binding; in plum, loop 6 stabilizing glutamate; and in light brown, the catalytic base. Underlined in gray, loop 6; underlined in plum, mutated loop 6 amino acids. Selected experimentally confirmed functional class I EAL domains with conserved signature amino acids (green bar); functional class IIa and IIb EAL domains with partially deviating signature amino acids (red bar), and nonfunctional class IIIa and IIIb EAL domains (blue bar) have been used in the alignment. Figure 1c. Phylogenetic tree of EAL domains of proteins most similar to ZMO1055_zm4_. Green: all conserved signature amino acids with demonstrated functionality are conserved. Blue: some conserved signature amino acids with demonstrated functionality are not conserved. Proteins used for the alignment and tree construction are indicated in Supplementary data.

Proteins ZMO0401, ZMO1487 and ZMO1055 contain an EAL domain which conventionally displays PDE activity. Possessing a conservative and a non-conservative amino acid exchange in the signature motifs, residue Y341 had been substituted by phenylalanine (F) in ZMO0401 and ZMO1487 and N417 substituted by methionine (M) in ZMO1487, respectively. With all other signature amino acid motifs present, including the catalytic base glutamate in the EGxE motif, a PDE activity has been demonstrated for ZMO0401 and ZMO1487. The signature amino acid motifs required for catalytic activity of the EAL domain of ZMO1055 are conserved as expected from its apparent PDE activity (Figure 1b and S1b; (Li et al., 2023a)).

However, our previous result indicated that a C1577T mutation in the ZMO1055 open reading frame resulting in an A526V substitution which inhibited the apparent PDE activity (Cao et al., 2022). We were therefore wondering about the impact of A526 in EAL domains. As a first step, with the GGDEF-EAL domain proteins MorA (PA4601) and PA0575 of *Pseudomonas aeruginosa*, the highest scoring templates for structural EAL domain models in a Phyre2 analysis as reference sequence, we analyzed the conservation of the amino acid sequences in the EAL domains in ZM4. As described above, the alignment of all *Z. mobilis* EAL domain showed that the vast majority of signature amino acid are conserved in the template and *Z. mobilis* EAL domains. Note that although the amino acid sequence identity is only 40.1% (similarity 56.7%) alanine at position 526 is conserved in MorA, but not in PA0575. However, a specific function has not been assigned to the equivalent of the 526 position. In contrast, in other *Z. mobilis* EAL domains, the amino acid A526 is not conserved (Figure S1b; (Rao, Qi et al., 2009)).

### Initial characterisation of basic catalytic features of ZMO1055

Complex GGDEF-EAL containing cyclic di-GMP turnover proteins, although catalytically competent, can affect phenotypes through a scaffold function independent of the catalytic activity, by targeted diguanylate cyclase/phosphodiesterase activity or alternative catalytic activity. Such an example is YciR, a GGDEF-EAL domain protein of *Escherichia coli* and *Salmonella typhimurium*, which modulates expression of the rdar biofilm activator *csgD* and subsequently rdar biofilm formation by interaction with alternative diguanylate cyclases and transcriptional regulators; and potentially alternative catalytic activity, respectively (Ahmad, Cimdins et al., 2017, Li, Yin et al., 2023b, Lindenberg, Klauck et al., 2013).

To assess the impact of amino acid substitutions on the phosphodiesterase activity in *Z. mobilis* ZM401, we first set up the phenotypic assays. In this context, we investigated whether ZMO1055 exerts its effect on flocculation in *Z. mobilis* ZM401 due to its phosphodiesterase activity (Li et al., 2023a). We chose alanine substitution of E356 in the conserved EAL motif with the glutamate involved in divalent ion binding and of the catalytic base E536 in the E_536_GVE conserved motif required for catalytic activity. These mutants also served as a negative control to assess potential differences in the apparent DGC and PDE activity of ZMO1055 between ZM401 and ZM4.

As a reference phenotype, upon expression of ZMO1055_ZM4_ in the constitutively flocculating strain *Z. mobilis* ZM401, the self-flocculation phenotype was completely disrupted as compared with the vector control (Figure 2; (Li et al., 2023a)). Expression of ZMO1055_ZM4_ variants E356A and E536A, however, did not alter the self-flocculation phenotype of strain *Z. mobilis* ZM401, indicating that both E356 and E536 are essential for phosphodiesterase activity of ZMO1055_ZM4_ and the PDE activity of the EAL domain is required to dissolve self-flocculation (Figure 2a). *Z. mobilis* ZM401 does not only show self-flocculation, but also displays a characteristic dye binding morphotype on Congo red (Figure 2b) and Calcofluor agar plates (Figure 2c). Mutant analysis showed that this dye binding morphotype, as the self-flocculation, is due to expression of the exopolysaccharide cellulose (Figure 2b and 2c and data not shown). The appearance of the colony morphology was included as an alternative assessment of the catalytic activity of ZMO1055_ZM4_ and its mutants. Production of ZMO1055_ZM4_ showed downregulation of the characteristic red rdar colony and downregulation of Calcofluor binding, both indicating that cellulose biosynthesis was dramatically downregulated. On the other hand, the enhanced reddish colony morphotype on the Congo red agar plate and enhanced fluorescence on the Calcofluor plate of strain *Z. mobilis* ZM401 expressing the ZMO1055_ZM4_ catalytic mutants in the EAL domain, E356A and E536A, suggested an even higher accumulation of cellulose triggered by the DGC activity of ZMO1055 (Figure 2b and 2c; (Li et al., 2023a)). Of note, upon overexpression of ZMO1055_ZM4_, a brown Congo red binding phenotype appeared temporarily within the first 24 h (Figure S2a). The molecular basis of this characteristic dye binding is currently unknown, but might be coupled to the DGC activity of ZMO1055_ZM4_ (Li et al., 2023a). These observations thus indicate a temporal switch between DGC and PDE catalytic activity as observed for other GGDEF-EAL proteins.

**Figure 2.**
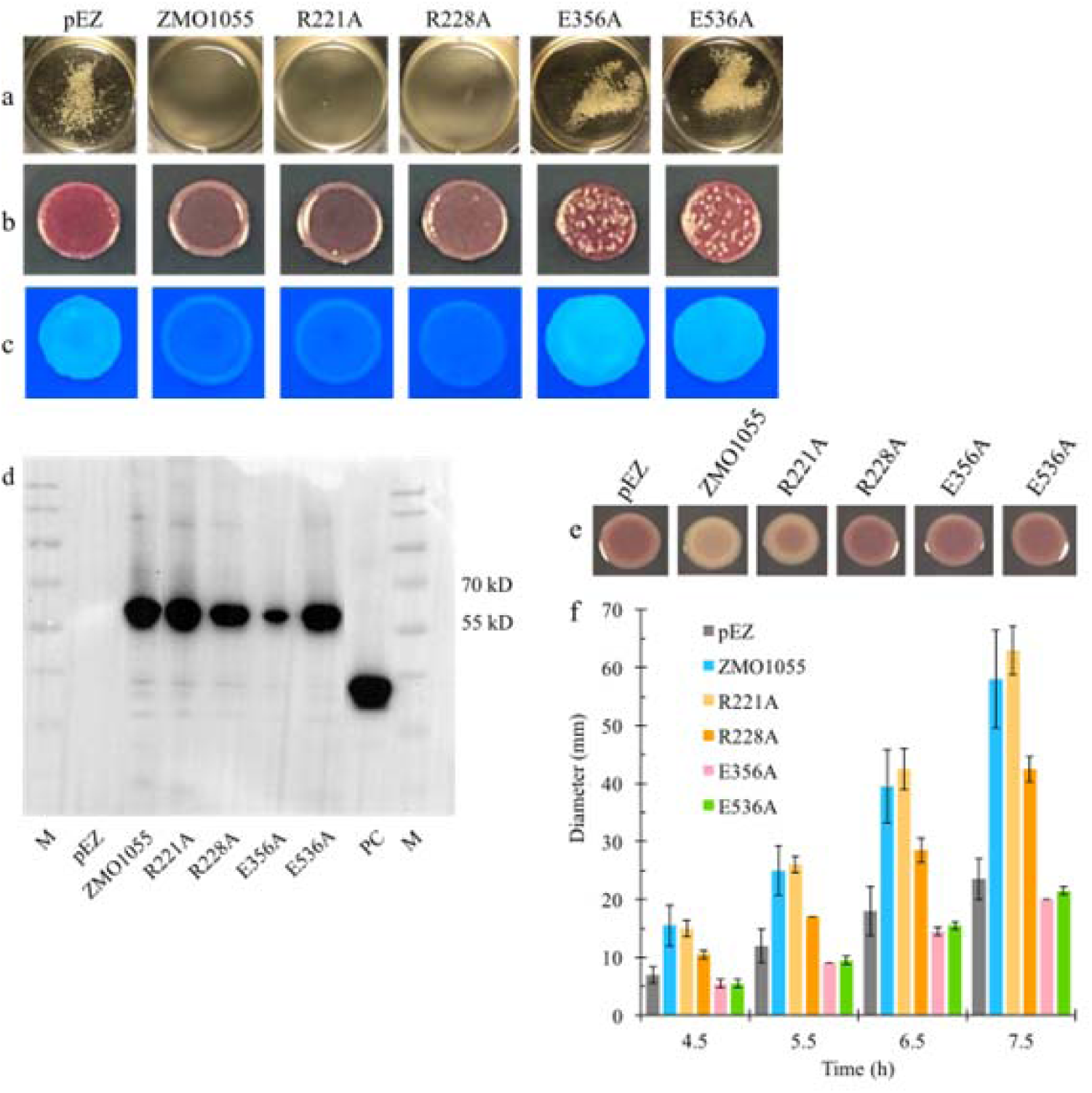
The PDE activity of ZMO1055 from *Z. mobilis* ZM4 is required for downregulation of flocculation in *Z. mobilis* ZM401 and up- and downregulation of motility and biofilm formation in *S. typhimurium* UMR1 Δ*yhjH*. Flocculation in liquid medium (a), Congo red assay (b) and Calcofluor White Staining assay (c) of *Z. mobilis* ZM401 strains over-expressing wild-type ZMO1055_ZM4_ or its DGC/PDE inactivated mutants after 48 and 24 h. Colony morphology morphotype of *Z. mobilis* ZM401 upon deletion of the cellulose synthease gene *bcsA* (d). Protein expression of ZMO1055_ZM4_ and mutants (e), CapV with 6xHis-tag from *E. coli* MG1655 (Li et al., 2022) was applied as a positive control for Western blotting. The rdar colony morphology assay (f) and swimming motility (g) of the *S. typhimurium* UMR1 Δ*yhjH* strain expressing wild-type ZMO1055_ZM4_ or DGC/PDE inactivated mutants.

As the GGDEF domain can also function as a sensory domain to affect the catalytic activity of the downstream EAL domain (Christen, Christen et al., 2005), we subsequently also assessed whether the PDE activity of ZMO1055_ZM4_ is affected by mutations in the GGDEF domain. We did not investigate the GGAQF catalytic mutant as it does not possess a phenotype in our assay (Li et al., 2023a). To this end, we chose to substitute R221 of the RxxD (I-site) motif, part of the cyclic di-GMP binding I-site (Christen, Christen et al., 2006) and R228 of the RxGGDEF conserved motif by alanine. Mutations R221A and R228A introduced into ZMO1055_ZM4_ caused the de-flocculation phenotype however retained few flocs. These data indicated that these mutations in the GGDEF domain either affect the catalytic activity of the GGDEF or EAL domain of ZMO1055_ZM4_, but only to a minor extent. The impairment of the allosteric I-site by the R221A substitution might enhance the DGC activity of the GGDEF domain of ZMO1055_ZM4_ (Christen et al., 2006, Li et al., 2023a).

To evaluate whether the observed changes in the self-flocculation morphology are due to differences in protein expression level or protein activity, relative expression of ZMO1055_ZM4_ and mutants was assessed by Western blot analysis using 6xHis-tagged proteins. The results showed that production of the ZMO1055_ZM4_ variants with R228A and E356A substitutions was impaired, but there was no correlation between expression level and self-flocculation (Figure 2d). Therefore, the difference in flocculation morphology are due to the amino acid mutation to affect the PDE activity thus causing no alteration, or even an increase in cellulose biosynthesis and self-flocculation compared to the vector control.

### *Heterologous expression of ZMO1055_ZM4_ and its mutants affect phenotypes in the* Salmonella typhimurium *UMR1 motility/sessility model*

The interaction of the ZMO1055 scaffold with other proteins in *Z. mobilis* ZM4 might alter the catalytic activity of the EAL domain. In order to assess the apparent catalytic activity of ZMO1055_ZM4_ and its mutants in a heterologous host where intermolecular protein-protein interactions most likely are substantially different, we choose as an alternative model to assess the effect of ZMO1055_ZM4_ and variants on expression of the rdar biofilm morphotype and flagella-based swimming motility of *S. typhimurium* UMR1 (ATCC 14028 Nal^r^ rdar_28°C_). To this end, the strains with a deletion in the phosphodiesterase required mainly for motility regulation, *S. typhimurium* UMR1 Δ*yhjH*, has been shown to be an ideal model to simultaneously assess PDE activity by downregulation of rdar biofilm morphotype expression and upregulation of flagella-based swimming motility (El Mouali et al., 2017). As addition of a 6x-His-tag can alter the functionality or degree of regulation of a protein, we first tested the effect of 6x-His on ZMO1055_ZM4_ and ZMO1055_ZM401_. Indeed, addition of 6x-His reduced the apparent phosphodiesterase activity of ZMO1055, but did not blur the effect of the A526V mutation (Figure S2b) Subsequently, we then assessed the catalytic mutants E356A and E536A in the EAL domain of ZMO1055_ZM4_. Those mutants did not alter the wild type phenotype with respect to rdar morphotype expression indicative for the loss of PDE activity of ZMO1055_ZM4_. On the other hand, swimming motility was even more repressed in these mutants indicative for a residual apparent DGC functionality of the mutant proteins (Figure 2e and 2f; (Li et al., 2023a)). Thus, the *Z. mobilis* and *S. typhimurium* based assays possess a different sensitivity to monitor the DGC activity of ZMO1055_ZM4_.

Upon expression of the ZMO1055_ZM4_ GGDEF domain mutants in strain *S. typhimurium* Δ*yhjH,* expression of wild-type ZMO1055_ZM4_ and its R221A mutant abolished the rdar phenotype. In congruency, these two strains displayed enhanced swimming motility, indicating that the mutations in the GGDEF domain of ZMO1055_ZM4_ had a minor effect on the apparent phosphodiesterase activity. However, even though the R228A mutant in the GGDEF domain maintained rdar phenotype expression as the vector control strain, it still displayed significantly enhanced swimming motility (Figure 2e and f). These observations indicate again a differential sensitivity of the two biological assays in the same strain for apparent catalytic activities. This observation is not surprising as cyclic di-GMP turnover proteins have been shown to act locally (Giacalone, Smith et al., 2018, Romling et al., 2013) and can be dedicated to distinct physiological processes such as the phosphodiesterase YhjH dedicated to flagella-based motility (Le Guyon, Simm et al., 2015). Alternatively, the R228A mutation has a complex effect on the apparent catalytic activity of ZMO1055_ZM4_ in the *S. typhimurium* model.

### The PAS-GGDEF-EAL domain structure is required for ZMO1055_ZM4_ functionality

Based on conserved domain analysis by alignment of homologous proteins and protein structural modeling by Phyre2 (Kelley et al., 2015, Liu, Lee et al., 2020), we concluded that ZMO1055 contains an N-terminal PAS (Per-Arnt-Sim) domain (Xing, Gumerov et al., 2023). PAS domains display a high sequence diversity explaining why the PAS domain of ZMO1055 has not been recognized by standard two dimensional BLAST search (Altschul, Gish et al., 1990). A PAS domain does not only function as a signal receiver and transducer domain, but dimerizes GGDEF/EAL domain containing proteins enabling or enhancing the catalytic activity upon signal perception (Schirmer, 2016).

We assessed the impact of domain interactions for the activity of ZMO1055_ZM4_ by selective domain overexpression. As reported above, upon overexpression of full length ZMO1055_ZM4_ in the self-flocculating strain ZM401, the self-flocculation phenotype of ZM401 was completely disrupted, concomitant with substantial degradation of intracellular cyclic di-GMP by ZMO1055_ZM4_ (Cao et al., 2022, Li et al., 2023a). We subsequently constructed different ZMO1055_ZM4_ variants with truncations of one or two domains. Overexpression of a ZMO1055_ZM4_ derived construct containing only the EAL domain did not lead to de-flocculate *Z. mobilis* ZMO401, indicating the necessity of the PAS and/or GGDEF domain in assisting PDE activity, dimerization and/or expression of ZMO1055_ZM4-EAL_ (Figure 3a). Alternative phenotypic assays on Congo red and Calcofluor plates indicated residual activity of ZMO1055_ZM4-EAL_ compared with the vector control (Figure 3b and 3c). Western blot analysis demonstrated that expression of the stand-alone ZMO1055_ZM4-EAL_ was not detectable in this assay, inidcating the requisiteness of the PAS and/or GGDEF domain or immediate adjacent amino acids required for expression of EAL domain.(Figure 3d).

**Figure 3.**
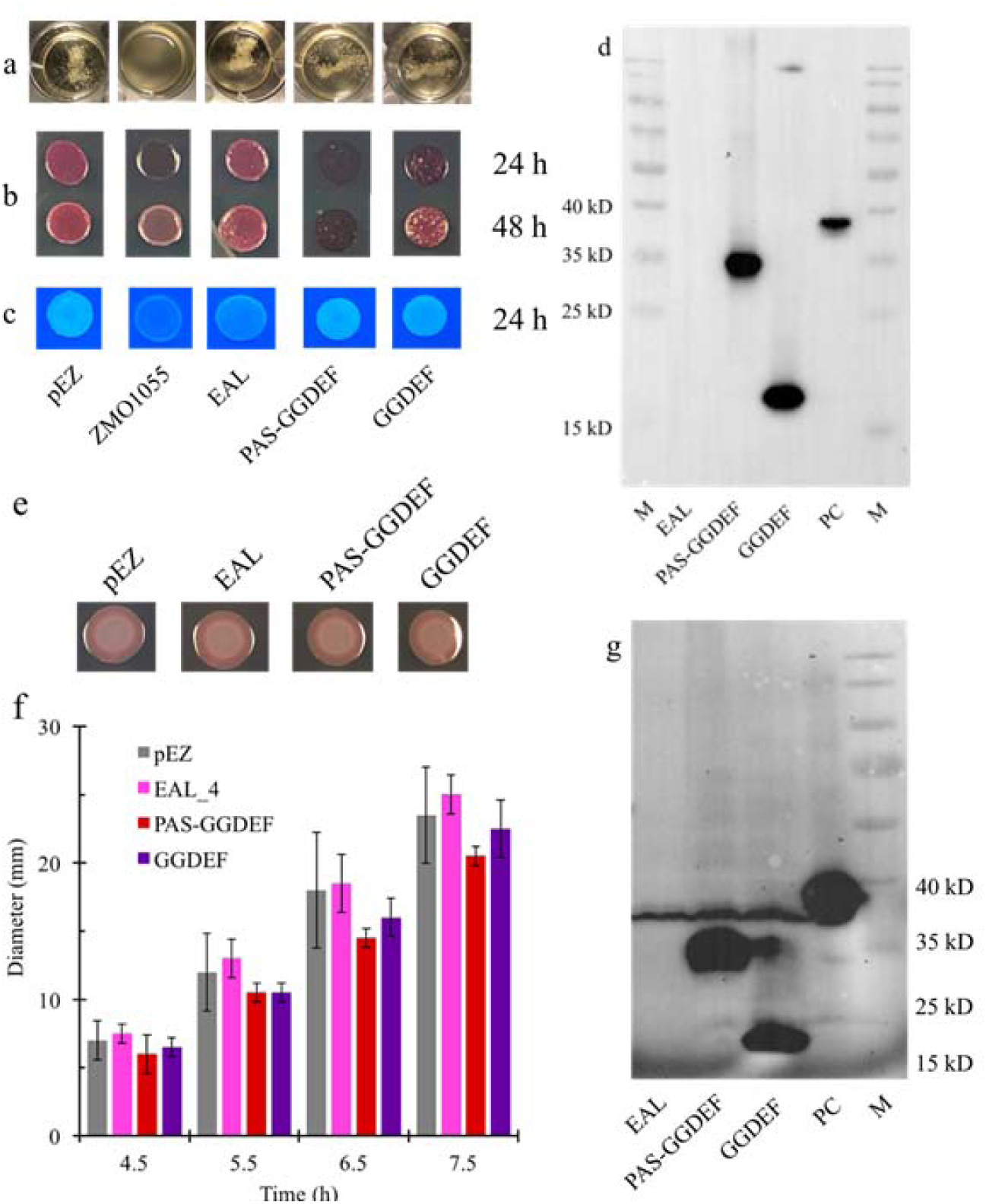
ZMO1055 of *Z. mobilis* ZM4 is a PAS-GGDEF-EAL domain protein with the full length domain structure required for activity. Flocculation (a), rdar morphotype (b) and Calcofluor White staining (c) assays of *Z. mobilis* ZM401 with overexpression of wild-type protein ZMO1055_ZM4_ and variants truncated in the individual domains. Production of the EAL, PAS-GGDEF and GGDEF domain in *Z. motilis* ZM401 with the 6x-His-tagged patatin-like phospholipase CapV as a positive control (d). Rdar assI(e), motility (f) and protein production (g) of *S. typhimurium* Δ*yhjH* producing the EAL, PAS-GGDEF and GGDEF domain of ZMO1055.

We further investigated whether the GGDEF domain has a function on its own. Overexpression of either PAS-GGDEF or GGDEF did not change the self-flocculation phenotype compared with the vector control, suggesting that ZMO1055_ZM4-GGDEF_ and ZMO1055_ZM4-PAS-GGDEF_ has no function on its own but requires the EAL domain for functionality in the self-flocculation assay. However, the Congo red assay exhibited a substantial alteration in colony morphology and inhibition of cell growth by overexpression of the PAS-GGDEF and GGDEF domain. Western blot analysis showed high expression of both constructs suggesting residual catalytic activity and/or GTP substrate binding of the GGDEF domain (Figure 3d (Li et al., 2023a)).

Assessment of the functionality and expression of the individual domains in *S. typhimurium* Δ*yhjH* as alternative and heterologous host showed no difference in rdar or motility morphology from vector control upon overexpression of the three constructs (Figure 3e and 3f). Thus, the truncated constructs are nearly inactive despite high expression of the PAS-GGDEF and GGDEF domain in the *Salmonella* system (Figure 3g). Again, ZMO1055_ZM4-EAL_ was not expressed consistent with the production experiments in *Z. mobilis* ZM401.

### Amino acids at position 526 differentially affect the apparent catalytic activity of ZMO1055

The amino acid sequence similarity among GGDEF/EAL domains encoded by one genome is usually less than 35% with *Z. mobilis* ZM4 no exception from other bacterial species. Such shows the EAL domain of ZMO1055 33.1 and 25.6% identity to the EAL domain of ZMO401 and ZMO1487. Thus we considered the possibility that A526, although not conserved among *Z. mobilis* ZM4 EAL domains has a specific role in one or more subgroups of EAL domains. We therefore analyzed amino acid conservation at the equivalent of the position 526 of the EAL domain in ZMO1055_ZM4_ in the first 5039 non-redundant most closely related proteins as derived from the NCBI database (Altschul et al., 1990). After automatic and subsequently manual curation of the alignment of the EAL domains of these proteins, a statistical summary of the conservation at this site was performed. Surprisingly, this analysis showed that upon analysis of 5039 sequences A536 is highly conserved among the ZMO1055 based subclass of GGDEF-EAL domain proteins present in over 70.5% of the sequences, while valine at this position was present at a frequency of 0.14% (Figure S3a). Other amino acids present were for example cysteine, with a frequency of 0.8%. The phylogenetic relatedness of the EAL domain of ZMO1055 was assessed by alignment with representative EAL domains. The phylogenetic tree constructed from the alignment in Figure 1c indicates that the EAL domain of ZMO1055_ZM4_, although closely related, forms an independent subclass distinct from other EAL domains.

This and previous studies had shown that the A526V mutation in ZMO1055_ZM401_ maintains cell aggregation and the intracellular cyclic di-GMP concentrations to a greater extent (Cao et al., 2022, Li et al., 2023a). The conservation of alanine at position 526 in subclass ZMO1055 GGDEF-EAL domain proteins indicates that alanine is important for functionality of the EAL domain. Thus, we raised the question how replacement of the equivalent position of alanine 526 by alternative amino acids affects the apparent catalytic activity of the EAL domain.

*Z. mobilis* ZM401 derived from *Z. mobilis* ZM4 by nitrosoguanidine mutation (Lee, 1982). Genomic comparison and molecular manipulation confirmed that the single nucleotide mutation C1577T in *ZMO1055_ZM4_* which resulted in A526V to be one of the two key events to trigger self-flocculation (Cao et al., 2022, Li et al., 2023a). The A526V substitution in ZMO1055 causes a distinct alteration in functionality.

Alanine is the amino acid with the shortest non-polar side chain of only one -CH_3_ group. The side chain of valine contains two additional non-polar -CH_3_ groups. Based on the structural model and molecular docking simulations we have hypothesized that the increased size of the side chain of valine affects the catalytic center and cyclic di-GMP binding (Cao et al., 2022).

To extend this observation and to gain insights into the molecular mechanism of the decreased apparent phosphodiesterase activity, we consequently we introduced amino acid substitutions by site directed mutagenesis at the position 526 to investigate their effect on the PDE activity of ZMO1055_ZM4_ Our choice was based 1. on the frequency of occurrence of alternative amino acids in the first 5039 homologs of ZMO1055 (Figure 3a); and 2. on amino acids with similar chemical properties, but larger size of the side chain compared with alanine and valine. To this end, A526 had been replaced by serine (frequency 2.4% at the 526 position), threonine (0.54%), isoleucine (0.04%) and leucine (not present but side chain only with different branching compared to isoleucine). Serine and threonine are amino acids with a polar side chain, but larger size of the side chain when compared with alanine. Isoleucine and leucine possess a larger aliphatic side chain than alanine and valine. Alanine was also substituted by glycine, with 24.5% the second most frequent amino acid at position 526, which confers alpha helix destabilizing properties.

We then assessed the effect of the amino acid substitutions with ZMO1055 variant expression in the *Z. mobilis* ZM401 and *Salmonella* model systems with the proteins ZMO1055_ZM4_ and ZMO1055_ZM401_ as references. In *Z. mobilis* ZM401 expressing ZMO1055_ZM4_ and its variants, substitution of alanine by glycine, serine and threonine negatively affected the de-flocculation activity compared to ZMO1055_ZM4_, to a similar extent as the substitution by valine, with few flocs remaining indicating diminished apparent PDE activity. Of note, those amino acids either possess no side chain or a side chain in size up to valine. On the other hand, overexpression of the protein variants with a leucine and isoleucine substitution not only retained but intensified self-flocculation with tighter flocks and a clear supernatant exceeding flocculation of the vector control indicating highly reduced apparent PDE enzymatic activity and/or elevated DGC activity (Figure 4a). Isoleucine and leucine possess a larger aliphatic side chain. As the self-flocculation phenotype mediated by cellulose biosynthesis is associated with elevated concentration of intracellular c-di-GMP which is effectively degraded by ZMO1055_ZM4_ (Li, Cao et al., 2022, Li et al., 2023a), the size of the side chain of the 526^th^ amino acid significantly affects the apparent PDE activity most likely by steric hindrance with access of cyclic di-GMP to the active site impaired (Cao et al., 2022).

**Figure 4.**
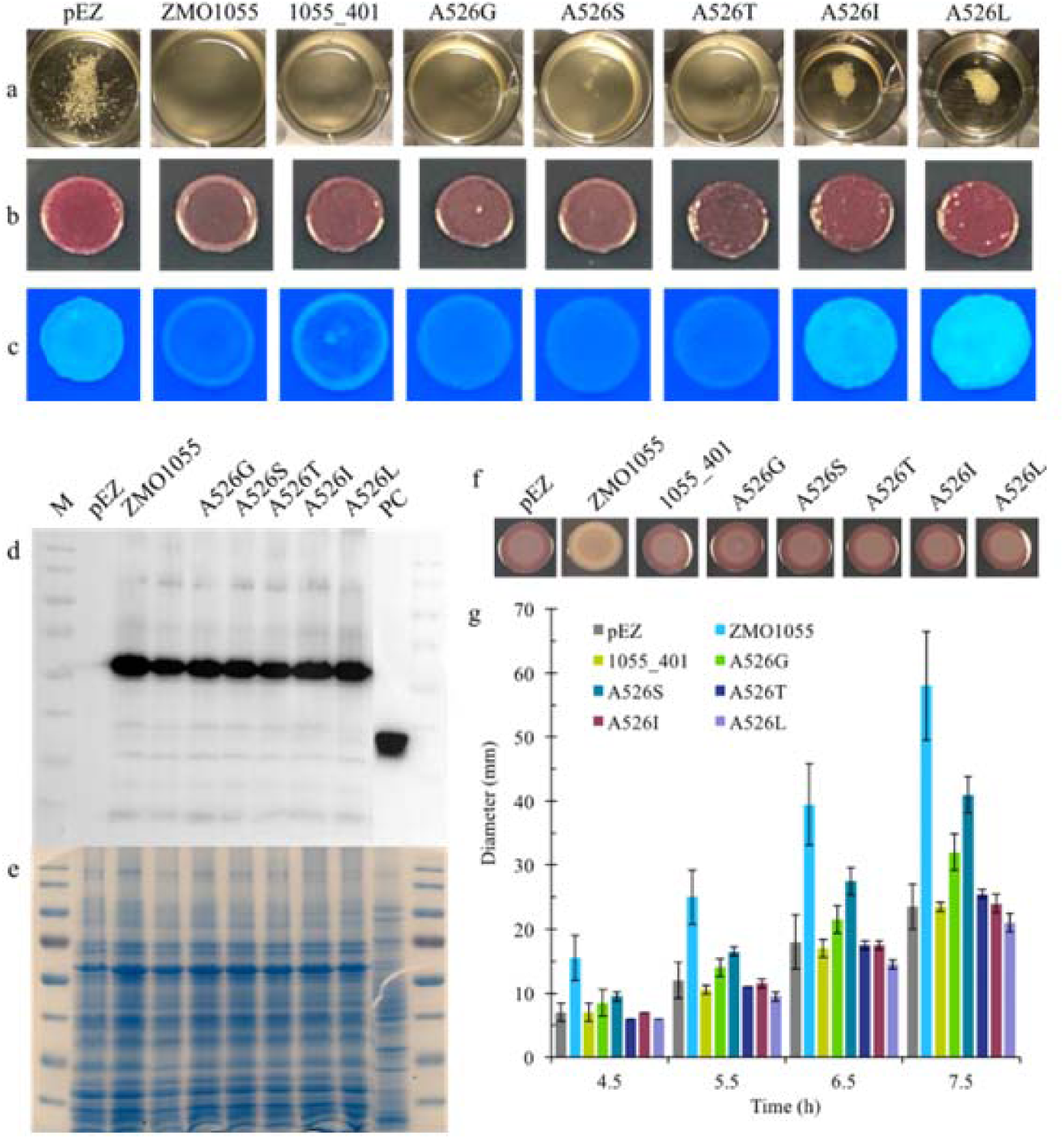
The size of the amino acid side chain at the A526 site of ZMO1055 regulates the apparent PDE activity. Flocculation (a), and rdar morphotype (at 48 h) and Calcofluor White (at 24 h) (b) assays of *Z. mobilis* ZM401 overexpressing ZMO1055_ZM4_ or the amino acid A526 substituted by valine, glycine, serine, threonine, isoleucine and leucine. Protein expression in *Z. mobilis* ZM401 was detected by western blot (c) and SDS-PAGE (d). Rdar phenotype (e) and swimming motility (f) of *S. typhimurium* UMR1 Δ*yhjH* strains expressing ZMO1055_ZM4_ or related protein variants.

Besides the self-flocculation phenotype, we assessed the effect of alterations in apparent PDE activity by monitoring cellulose production by the Congo red and Calcofluor white binding assay in *Z. mobilis* ZM401. The reference proteins ZMO1055_ZM4_ and ZMO1055_ZM4_ A526V (ZMO1055_ZM401_) upon overexpression showed a characteristic Calcofluor white binding patterns with dye binding at the rim of the colony only. This observation indicates that the phosphodiesterase activity of ZMO1055 is highly active in late phase cells, but inactive in exponentially growing cells consistent with the temporal observations (Figure S2a). In contrast, overexpression of ZMO1055_ZM4_ containing either the A526S, the A526T and the A526G substitution showed equal low level Calcofluor white binding throughout the colony indicating still effective apparent PDE activity. The Congo red binding pattern upon overexpression of all three ZMO1055_ZM4_ variants followed this trend. These observations confirmed the hypothesis that the amino acid at position 526 affects the apparent phosphodiesterase activity through steric hindrance rather than by the chemical property of the amino acid side chain. Further, the mutations caused growth phase independent apparent phosphodiesterase activity of ZMO1055.

In contrast, overexpression of ZMO1055_ZM4_ with the A526I or A526L substitutions to cause dark red morphotypes with rough surface and bright Calcofluor white staining higher than the vector control (Figure 4b, c). In conclusion, the Congo red and Calcofluor assay grossly confirmed the direction of the phenotypic consequences upon expression of the ZMO1055_ZM4_ mutants. In particular, the longer branched aliphatic side chains of the amino acids leucine and isoleucine in substitution for alanine at position 526 retained the *Z. mobilis* ZM401 phenotype or even enhanced the cellulose-based phenotypes cell aggregation and dye binding capacity. Although these findings confirmed our hypothesis that amino acid 526 affects PDE activity through steric hindrance rather than by chemical property, the effect of the A526G mutation still needs to be explained.

Western blot analysis was applied to assess the expression of ZMO1055_ZM4_ in comparison to its 526 position variants. When the total protein content based on assessment of the protein pattern after Coomassie staining for a 12% SDS-PAGE gel had been normalized and expression of ZMO1055_ZM4_ assessed, we concluded that production of variant proteins was not affected by the different mutations. Thus the different self-flocculation phenotypes were due to the altered apparent PDE activity of ZMO1055_ZM4_ variants rather than changed expression level (Figure 4c, d and e).

Heterologous expression of the mutant proteins in the model of the rdar phenotype of *S. typhimurium* Δ*yhjH* was indiscriminatory as a nearly similar colony phenotype without any difference in color or roughness as the vector control was observed (Figure 4f). On the other hand, swimming motility of *S. typhimurium* Δ*yhjH* strains showed a differentiated behavior of the mutants: elevated motility was observed upon expression of ZMO1055_ZM4_ with the A526S substitution and ZMO1055_ZM4_ with A536G. However, the increase in motility did not reach the ZMO1055_ZM4_ level. Comparable to ZMO1055_ZM401_ (ZMO1055_ZM4_ A526V), expression of ZMO1055_ZM4_ A526T, A526I and A526L did not elevate swimming motility over the vector control in *S. typhimurium* Δ*yhjH* under the applied experimental conditions (Figure 4g). Therefore, the apparent PDE activity of ZMO1055_ZM4_ in *S. typhimurium* is affected by the 526^th^ amino acid grossly depending on the size of the side chain.

### Structural model of the effect of amino acids predicted to be involved in steric hindrance in ZMO1055_ZM4_

A structural model of ZMO1055_ZM4_ was constructed by de novo calculation with Phyre2 and I-TASSER as well as with SWISS-MODEL with 5xgb.1.A from *Pseudomonas aeruginosa* as reference structure (Figure 5a and data not shown; (Kelley et al., 2015)). Similar structural characteristics such as the arrangement of secondary structures and locations of conserved sites were exhibited when comparing the three 3D structures of ZMO1055_ZM4_. Furthermore, amino acid 526 locates close to the end of an α-helix facing the active center of the EAL domain. However, the side chain of amino acid 526 does not establish direct contact with the active site but is separated by two β-strands formed by amino acids D497-I501 and I531-E536. To test whether the apparent PDE catalytic activity could be recovered in ZMO1055_ZM401_ via those two β-sheets, the amino acids L499 and I531 were chosen to be substituted by amino acids with shorter side chains as they are in the line from A526 to the active center based on the structural model (Figure 5a). Leucine and isoleucine constitute 96.3% of the amino acids present at position 499 and 45.1% at position 531 among the first 5039 homologs of ZMO1055.

**Figure 5.**
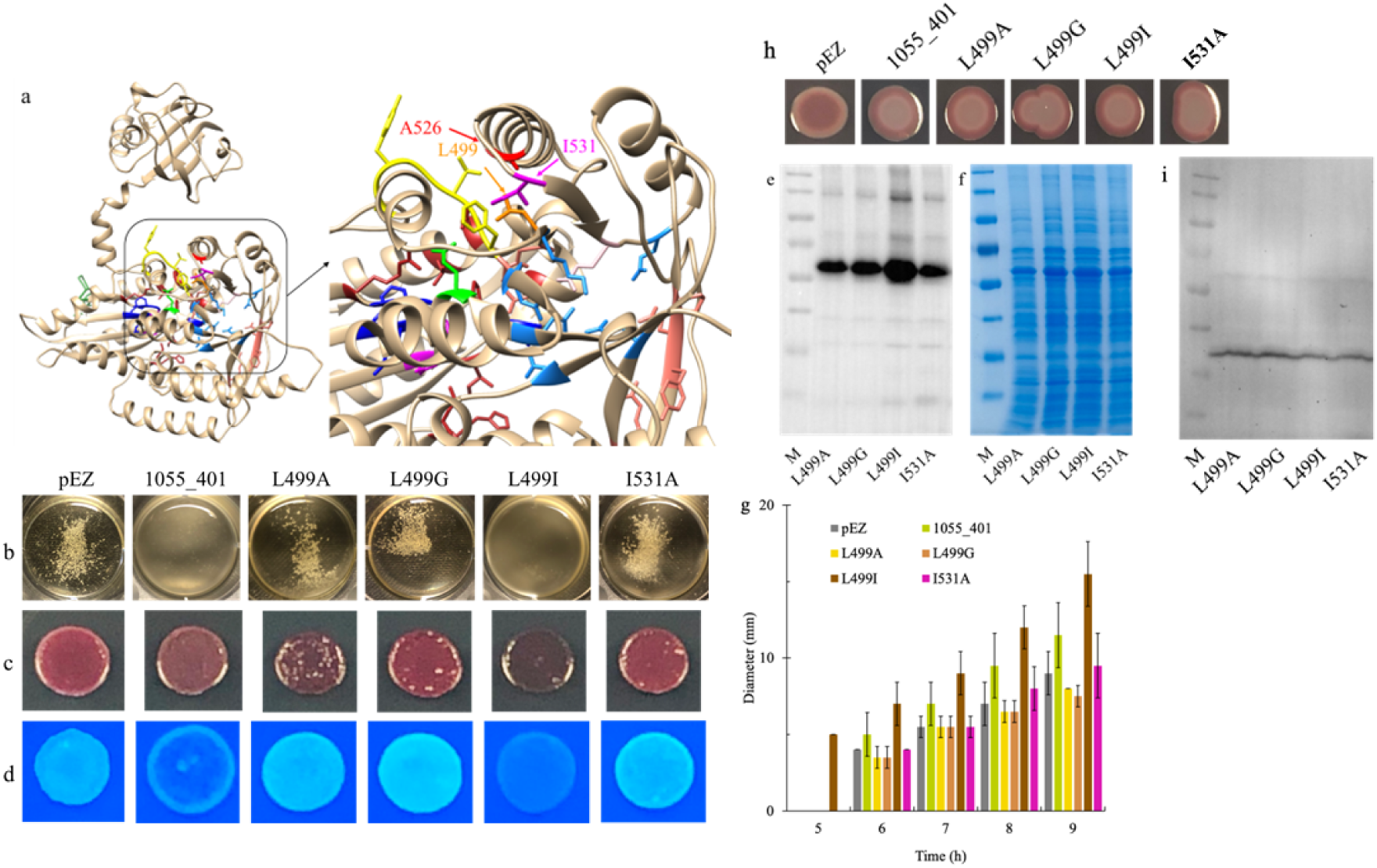
Possible mechanism applied by mutation site. Structural modeling of ZMO1055_ZM4_ by Phyre2 (a). A forest green color indicates the allosteric I-site, firebrick indicates the substrate interacting sites, blue indicates sites stabilizing the transition state, magenta indicates the Mg^2+^ binding sites in the GGDEF domain and yellow indicates Loop 6, green indicates Loop6 stabilizing sites, salmon indicates substrate binding sites, pink indicates the catalytic base, dodger blue indicates the Mg^2+^ binding sites, orange indicates L499 and magenta indicates I531 in the EAL domain. The mutation site (526^th^ amino acid) was colored in red and chain B of the structural model created by SWISS-MODEL was colored in sky blue. Flocculation (b), rdar (at 48 h) (c) and Calcofluor White staining (at 24 h) (d) assay of ZM401 mutants with overexpression of ZMO1055_ZM401_ and its L499A, L499G and I531A variant. Protein expression of ZMO1055_ZM401_ by Western blot (e) and SDS-PAGE (f). Swimming motility (g), rdar biofilm colony morphotype (h) and protein expression (i) of *S. typhimurium* Δ*yhjH* strains expressing ZMO1055_ZM401_ or respective protein variants.

We subsequently replaced L499 or I531 by alanine and glycine in ZMO1055_ZM401_, amino acids with smaller or no side chain. Only 0.1% of amino aicds at position 531 are alanine. In addition, we replaced L499 by its isomer isoleucine, which has the branching at the γ-carbon instead of the δ-carbon. Of note, we performed these experiments under condition where ZMO1055_ZM401_ (ZMO1055_ZM4_ A526V) showed a substantial de-flocculation phenotype in order to assess the extent of lower apparent PDE activity. The L499I mutation of ZMO1055_ZM401_ caused de-flocculation of *Z. mobilis* ZM401 cell aggregates similar as ZMO1055_ZM401_ (Figure 5b), while the Congo red and Calcoflour assay indicated higher apparent PDE activity. Overexpression of the ZMO1055_ZM401_ L499A and L499G variants equally as the I531A variant in *Z. mobilis* ZM401, however, surprisingly displayed flocculation exceeding the *Z. mobilis* ZM401 vector control as well as enhanced Calcofluor white binding indicating elevated cellulose production. Congo red binding was also enhanced, with distinct coloration upon overexpression of the two variants, Equally, the enhanced flocculation and Congo red and Calcofluor white dye binding capacity upon overexpression of the ZMO1055_ZM401_ I531A mutant compared to the vector control showed higher cellulose production and thereby the opposite than expected phenotypes (Figure 5c and 5d). Therefore, those two amino acids are involved in the regulation of the catalytic function of ZMO1055_ZM401_; most likely by highly downregulating the catalytic activity of the EAL domain thereby showing residual apparent diguanylate cyclase activity. The effect of those amino acid substitution on enzyme functionality is currently unpredictable by the structural model(s). We can however hypothesize that a distinct tight interaction between the aliphatic site chains of those proteins is required for catalytic activity, but lost upon the introduction of amino acids with other aliphatic side chains at the different positions in the protein.

Assessment of protein expression by western blot analysis indicated that all ZMO1055_ZM401_ mutants had equal expression levels than the parent protein ZMO1055_ZM401_ (Figure 5e, 5f, 4d, and 4e).

When ZMO1055_ZM401_ was heterologously expressed in *S. typhimurium* Δ*yhjH*, introduction of the ZMO1055_ZM401_ L499I mutant contributed to the stimulation of swimming motility to a higher extent than ZMO1055_ZM401_, while expression of the other double mutant constructs had no effect or even repressed swimming motility (Figure 5g). These results reflect the effect of those mutants upon apparent PDE activity in *Z. mobilis* ZM401. However, the rdar biofilm assay of *S. typhimurium* UMR1 Δ*yhjH* has been insensitive as it did not show any difference upon expression of the four proteins compared to ZMO1055_ZM401_ (Figure 5h). Protein production was not altered in *S. typhimurium* UMR1 Δ*yhjH* (Figure 5i).

### The effect of the A526V mutation is displayed also in other GGDEF-EAL domain proteins

To assess whether the effect of the A526V mutation on apparent PDE activity is conserved in EAL-domain containing proteins, we choose to investigate two closely related GGDEF-EAL proteins and three more distantly related EAL proteins which had an alanine at the same position as ZMO1055_ZM4_. According to phylogenetic analyses, two of the closely related proteins chosen were ARS29551.1 and WP_060851252.1, GGDEF-EAL proteins from *Sphingomonas* sp. KC8 and *Methylobacterium aquaticum* MA-22A, respectively (Figure 6a). A first protein with low amino acid similarity but with an alanine at the position equivalent to 526 is PA3258, an EAL-CBS-GGDEF protein obtained from *Pseudomonas aerginosa* SG17M. No phenotype had been detected for PA3258 in *P. aeruginosa* strains PAO1 and PA14 by transposon mutagenesis and gene over-expression (Kulesekara, Lee et al., 2006), while in *Pseudomonas fluorescens* Pf0-1 the homologous phosphodiesterase RapA depletes cyclic di-GMP upon phosphate starvation (Newell, Boyd et al., 2011). Two additional distantly related proteins with an alanine at a corresponding position were STM0468 and STM3615 from *Salmonella typhimurium* ATCC 14028 (for convenience the nomenclature of *S. typhimurium* LT2 is used, with the gene cloned from the clonal variant *S. typhimurium* UMR1 (ATCC 14028 Nal^r^ rdar_28°C_). To this end, the A to V amino acid substitutions were introduced at the site corresponding to A526 of ZMO1055_ZM4_ by site-directed mutagenesis.

**Figure 6.**
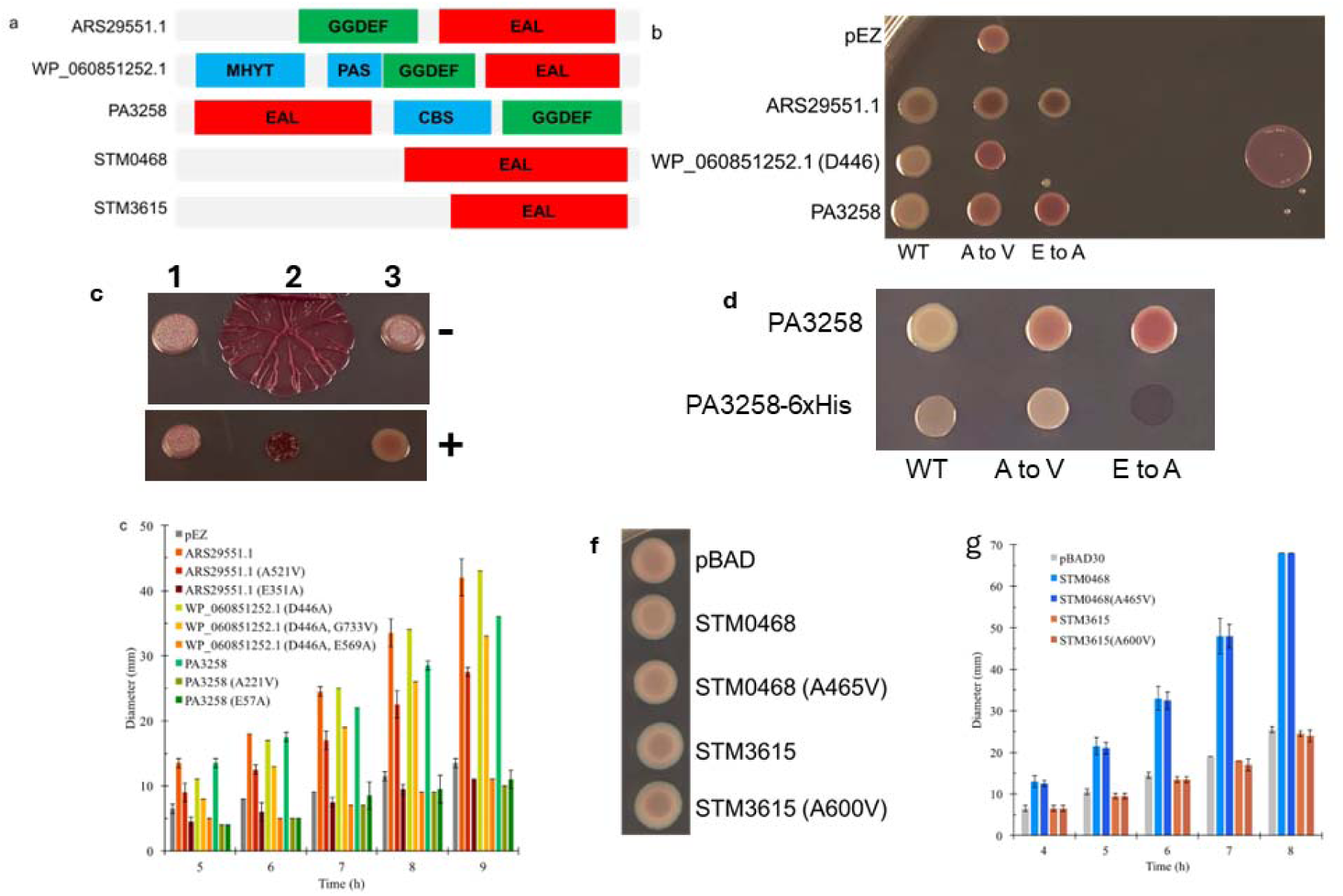
Effects of substitution of the equivalent of the A526 site on apparent PDE catalytic activity in GGDEF-EAL or EAL proteins closely and more distantly related to ZMO1055. Domain structure of the five investigated EAL proteins (a); Rdar colony morphology type (b) and flagella-based swimming motility (e) of *S. typhimurium* UMR1 Δ*yhjH* strains expressed with closely related GGDEF-EAL proteins. Rdar colony morphology type upon overexpression of WP_060851252.1 (D446A) versus WP_060851252.1 in *S. typhimurium* UMR1 Δ*yhjH* (c) upon induction by 100 ng/L anhydrotetracycline. *S. typhimurium* UMR1 Δ*yhjH*, 1. vector control pEZ; 2, WP_060851252.1 cloned in pEZ; 3, WP_060851252.1 D466A cloned in pEZ. Effect of a 6xHis-tag on the colony morphology type of PA3258 overexpressed in the host *S. typhimurium* UMR1 Δ*yhjH* (d). Rdar colony morphology type (f) and flagella-based swimming motility (g) of *S. typhimurium* UMR1Δ*yhjH* strains expressed with distantly related EAL proteins. (b, d, f) The rdar colony morphology type has been monitored after 24 h of growth at 28°C on Congo red agar plates. ( c) Flagella-based swimming motility has been monitored at indicated time points with motility LB 0.25% agar plates.

All the five gene products and their A to V substitution mutants were expressed in the *S. typhimurium* UMR1 Δ*yhjH* model strain to assess their effects on the apparent PDE catalytic activity (Figure 6b-e), although this type of phenotypic alteration can also be caused by overexpression of a cyclic di-GMP binding protein. As substantial apparent diguanylate cyclase activity was observed for WP_060851252.1 (Figure 6c), the DGC activity was abolished by mutating the GGDEF motif to GGAEF (WP_060851252.1 (D446)) to prevent interference with the assessment of the PDE activity. When the wild type ARS29551.1 protein and WP_060851252.1 (D446A) were expressed in the *S.* Typhimurium UMR1 Δ*yhjH* rdar biofilm model, the colonies appeared whitish compared with the pEZ vector control, confirming apparent PDE activity. However, when the A to V mutants were expressed, the color of the colony turned more reddish, suggesting reduced apparent PDE activity of ARS29551.1 and WP_060851252.1 (D446A) consistent with the predictions (Figure 6b). Negative controls that abolished the catalytic activity of the EAL domain were constructed by site-directed mutagenesis, replacing the glutamate by alanine in the EAL motif of ARS29551.1 and WP_060851252.1 (D446A). The rdar morphotype of ARS29551.1 with catalytically inactive EAL domain corresponded to the phenotype of the A521V mutant while production of WP_060851252.1 with the catalytically inactive EAL domain displayed a stronger rdar morphotype than its A724V mutant (Figure 6b).

Expression of the distantly related protein PA3258 C-terminally tagged with 6xHis in *S. typhimurium* Δ*yhjH* showed an extreme and incongruent morphotype (Figure 6c). We reasoned that the 6xHis-tag interferes with the protein functionality and/or stability. Upon expression of PA3258 without a 6xHis-tag a DGC (upon abolishment of the phosphodiesterase activity) and PDE positive phenotype was displayed (Figure 6b). As consistent to the other two proteins, the A221V mutation upregulated the rdar morphotype (Figure 6b). Furthermore, stimulation of motility was diminished for all A to V mutants compared to their PDE competent wild type counterparts, hinting substantial, but mostly incomplete inhibition of the apparent PDE activity in ARS29551.1 and WP_060851252.1 (D446A) (Figure 6d). Upon overexpression of the PDE inactivated mutants, motility of *S. typhimurium* UMR1 Δ*yhjH* was even downregulated compared to the vector control in all instances, indicating residual DGC activity of the proteins (Figure 6e).

The effect of an A to V substitution was also investigated in two other distantly related proteins, the EAL protein STM0468 and the GGDEF-EAL domain protein STM3615 from *S. typhimurium* (Simm, Lusch et al., 2007). An alanine is conserved in distantly related EAL domain proteins even without a GGDEF domain. Even though STM0468 showed PDE activity in the motility assay, its A465V mutation did not show visible downregulation of the apparent catalytic activity (Figure 6f and g). STM3615 and its A600V mutant had no effect on motility, demonstrating that STM3615 does not have a phenotype in the *S. typhimurium* Δ*yhjH* background as previously reported for the *S. typhimurium* UMR1 wild type (Figure 6f and g; (Anwar, Rouf et al., 2014, Simm et al., 2007)). Furthermore, no change in apparent activity was observed upon expression between the wild type and the mutant protein upon assessment of the rdar morphotype (Figure 6h). In conclusion, the effect of the A to V mutation on apparent PDE activity is conserved in GGDEF-EAL proteins ARS29551.1, WP_060851252.1 which belong to the ZMO-related clade and in distantly related PA3258, but not in STM0468 which both possess a distantly related EAL domain.

### Effect of amino acid 525 on the apparent phosphodiesterase activity of ZMO1055

Upon analysis of the conservation of alanine at the 526^th^ amino acid position in the aligned protein sequences, we noticed that only few mainly non-polar amino acids are present at the position 525 (Figure S3a). Methionine 525 as in ZMO1055 is present at a frequency of 31.8% in the first 5039 homologous proteins (Figure S3a). Further analysis on the first 5039 proteins most similar to ZMO1055 showed that the frequency of leucine is 64.14%. Therefore, methionine was substituted by leucine constructing a M525L mutant in ZMO1055_ZM4_ to explore its effect on cyclic di-GMP relevant phenotype.

Upon expression of ZMO1055_ZM4_ M525L in *Z. mobilis* ZM401, this protein variant diminished flocculation, rdar biofilm and Calcofluor white to a lesser extent than ZMO1055_ZM4_, but substantially more than ZMO1055_ZM401_, the ZMO1055_ZM4_ A526V mutant (Figure 7 a, b and c). Similarly, the M525L substitution in ZMO1055_ZM401,_ the ZMO1055_zm4_ A526V mutant, equally showed phenotypes consistent with a strong reduction of the apparent PDE activity compared to the ZMO1055_ZM401_ reference with substantial self-flocculation remaining. Thus, the M525L substitution combined with A526V reduced the apparent PDE activity of ZMO1055 substantially as flocs were still retained and the dye binding capacity with respect to Congo red and Calcofluor white has been substantially enhanced (Figure 7). Leucine has a non-polar branched side chain consisting only of methyl groups compared to methionine with a linear side chain. Therefore, as a hypothesis, the amino acid at 525^th^ site affects PDE activity by size and spatial arrangement like amino acids at the 526^th^ position.

**Figure 7.**
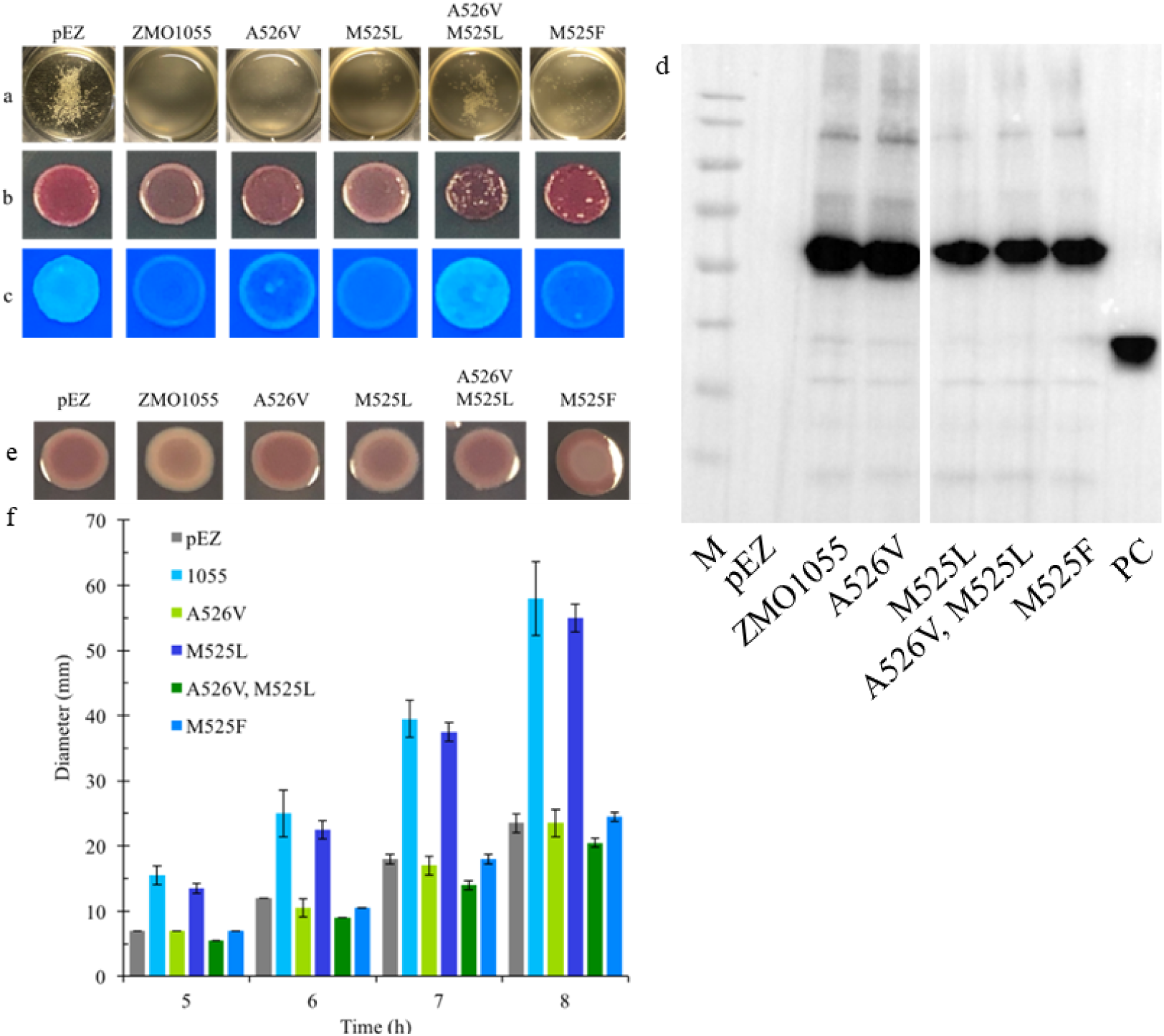
Effect of amino acid substitutions in the M_525_A motif on cyclic di-GMP relevant phenotypes. Flocculation (a), rdar morphotype (at 48 h) (b) and Calcofluor White staining (at 24 h) (c) assays of *Z. mobilis* ZM401 with overexpression of ZMO1055_ZM4_ or its M_525_A motif substitutes. Western blot demonstrating expression of ZMO1055_ZM401_ mutants in *Z. mobilis* ZM401 (d). Rdar colony morphology biofilm (e) and swimming motility (f) of *S. typhimurium* UMR1 Δ*yhjH* strains expressing ZMO1055_ZM401_ or its M_525_A motif substitutes. (e) The rdar colony morphology type has been monitored after 24 h of growth at 28°C on Congo red agar plates. (f) Flagella-based swimming motility has been monitored at indicated time points with motility LB 0.25% agar plates.

As the effect of the M525L substitutions are significant, but not substantial in all models, methionine at 525^th^ site was also substituted by phenylalanine, an amino acid present in 0.06% of the 5039 proteins most homologous to ZMO1055_ZM4_. Compared with ZMO1055_ZM4_ and ZMO1055_ZM4_ M525L, expression of ZMO1055_ZM4_ M525F in *Z. mobilis* ZM401 retained stronger self-flocculation, triggered a more pronouced rdar phenotype, but surprisingly highly reduced Calcofluor white staining (Figure 7 a, b and c). In conclusion, the amino acid at the 525^th^ position affects the activity of ZMO1055 context dependent with respect to the relevant cyclic di-GMP regulated phenotypes with the molecular mechanism still to be explained.

The *S. typhimurium* Δ*yhjH* model was subsequently chosen to further assess the effect of the methionine 525 substitution. A similar trend with a diminished apparent PDE activity for the M525L mutation in ZMO1055_ZM4_ and ZMO1055_ZM401_ as upon expression in *Z. mobilis* ZM401 was exhibited in *S. typhimurium* UMR1 Δ*yhjH* (Figure 7 d and e). The M525L mutant in ZMO1055_ZM4_ and ZMO1055_ZM401_ even slightly reduced swimming motility compared to the parent proteins. Expression of ZMO1055_ZM4_ M525F even enhanced rdar morphotype expression compared to the vector control and had no effect on motility as expression of ZMO1055_ZM401_ (comparable to the vector control). These results showed that the ZMO1055_ZM4_ M525F mutant displayed an apparently highly impaired PDE activity in *S. typhimurium* Δ*yhjH* in contrast to *Z. mobilis* ZM401. In conclusion, substituting amino acid not only at the 526^th^, but also at the 525^th^ position affects the apparent PDE activity of ZMO1055 with the M525F substitution to exceptionally show divergent phenotypic effects in *Z. mobilis* ZM401 and *S. typhimurium* UMR1 Δ*yhjH*.

## **D**iscussion

In this work, we investigated the impact of the A526V substitution in the PAS-GGDEF-EAL domain protein ZMO1055 which caused a significant downregulation of the apparent catalytic activity of its EAL domain phosphodiesterase associated with enhanced cellulose production and flocculation in *Z. mobilis* ZM4 (Cao et al., 2022, Li et al., 2023a). Our major findings include the observations: that

1, A526 is prevalent in the ZMO1055 subfamily of EAL domains, while valine at this position is infrequent;
2. replacement of A526 by amino acids with a more bulky side chain gradually cause downregulation of apparent phosphodiesterase activity most likely by steric hindrance of access of substrate;
3. the effect of the A526V and other substitutions is conserved in different model systems
4. the effect of the A526V and other substitutions is conserved in subgroups of EAL domains;
5. the size of the side chain of the amino acid at position 525 has a similar effect on the apparent phosphodiesterase activity.

Overall this study shows the delicate regulation of cyclic di-GMP signaling and its physiological output by even only a single conserved amino acid substitution in the EAL domain.

Although non-synonymous single nucleotide polymorphisms occur in open reading frames of bacterial isolates, clones and, more frequently among bacterial clones, their impact on protein functionality and subsequently microbial physiology and metabolism has been rarely investigated, perhaps due to the lack of screenable output phenotypes. It has been observed thought that single or multiple amino acid substitutions in cyclic di-GMP turnover proteins outside of the conserved signature motifs can have a profound impact on readily screenable phenotypes like colony biofilm types potentially through alteration of the catalytic activity of the domain, alterations of protein-protein interactions, alteration of signal perception or other mechanisms (Beyhan &Yildiz, 2007, Cimdins-Ahne, Naemi et al., 2023, Cimdins et al., 2017). However, although single or multiple amino acid substitutions of cyclic di-GMP turnover proteins are frequently observed in databases, effects of amino acid substitutions outside the conserved signature motifs are currently hardly predictable but requires experimental evidence for their effect on relevant phenotypes.

In this context, amino acids with aliphatic side chains can be highly conserved in proteins and provide a functionality beyond a plain structural role. Although amino acids with aliphatic side chains have been shown to be involved in the high affinity binding of receptors to ligands such as cyclic di-GMP (Roelofs, Jones et al., 2015, Wang, Chin et al., 2016) and the functionality of the CsgD biofilm transcriptional regulator (Wen, Ouyang et al., 2017), the impact of the choice of the different aliphatic side chains has been rarely explored (Baumann &Zerbe, 2024). In this work, we demonstrate that the aliphatic site chains that differ by only one or two methyl groups and not directly located in the catalytic site can have a profound effect on the performance of the GGDEF-EAL diguanylate cyclase/phosphodiesterase ZMO1055. We hypothesize though, that a more determinative role of amino acids with aliphatic side chains extends beyond ZMO1055 to other cyclic di-GMP turnover proteins.

The phenotype to monitor the catalytic activity of ZMO1055 has been in the first hand self-flocculation of *Z. mobilis* ZM4 caused by secretion of cellulose. The exopolysaccharide cellulose, a 1,4-beta-D-glucan, is a (biofilm) extracellular matrix component of microbes throughout the phylogenetic tree mediating cell-substrate interactions such as adhesion under flow conditions and cell-cell interactions such as microcolony and biofilm formation and cell aggregation (flocculation) (Grantcharova, Peters et al., 2010, Jeon et al., 2012, Romling &Galperin, 2015, Xia, Liu et al., 2018). There exist at least five distinct cellulose biosynthesis gene clusters that can be discriminated by distinct accessory genes in the cluster (Romling &Galperin, 2015). As a major regulatory mechanism, cyclic di-GMP signaling post-transcriptionally activates cellulose biosynthesis via the C-terminal PilZ domain (Morgan, McNamara et al., 2014, Ryjenkov, Simm et al., 2006). However cyclic di-GMP signaling is not only tightly coupled to cellulose biosynthesis, but ubiquitously stimulates multicellular behavior in bacteria. It is thus expected that the elevated cyclic di-GMP levels caused by the diminished apparent phosphodiesterase activity of ZMO1055_ZM401_ also affect other physiological traits besides cellulose biosynthesis.

Thus not only flocculation based on cellulose production, but also the colony morphotype as a plate-grown biofilm model of *Z. mobilis* ZM401 and *S. typhimurium* UMR1 is a sensitive biological assay to assess the activity of cyclic di-GMP turnover proteins (Simm, Ahmad et al., 2014). To our current knowledge, although in *Z. mobilis* ZM401, self-flocculation and the colony morphotype is based on solely cellulose biosynthesis. while in *S. typhimurium* UMR1, expression of the biofilm transcriptional regulator CsgD activates cellulose biosynthesis and amyloid fimbriae to commonly produce the cyclic di-GMP dependent rdar morphotype, a diversity of colony morphotypes has been observed. This variability in morphotypes is reflected by the differential effect upon overexpression of ZMO1055 variants with the mostly conserved amino acids substitutions at the 525/526 site previously not implicated in functionality. On the other hand, flagella-regulated swimming motility repressed by cyclic di GMP signaling is another sensitive biological assay to assess the activity of cyclic di-GMP turnover proteins (Ryjenkov et al., 2006).

In the different bacterial species, cellulose biosynthesis is regulated by cyclic di-GMP turnover proteins with diverse domain composition and cellular localization which can be polar, dispersed, membrane-associated or cytosolic (Romling &Galperin, 2015, Tal, Wong et al., 1998). Besides membrane-associated enzymes, frequently, PAS-GGDEF or PAS-GGDEF-EAL domain proteins provide the cyclic di-GMP for the activation of cellulose biosynthesis (Liu et al., 2020). Thereby, the degree of activation of cellulose biosynthesis is not necessarily directly correlated with the cellular cyclic di-GMP levels. The lack of correlation between the actual cyclic di-GMP concentration and activation of cellulose biosynthesis by distinct enzymes shows the complex regulation of biochemical processes by second messenger signaling (Li et al., 2023a, Tal et al., 1998). Local (close association of the diguanylate cyclase with the cellulose biosynthesis complex (Abidi, Torres-Sanchez et al., 2022, Kader, Simm et al., 2006, Massie, Reynolds et al., 2012)) and global signaling events might play a role. In the case of ZMO1055, although the phosphodiesterase activity of the PAS-GGDEF-EAL domain protein is dominant in the *Z. mobilis* flocculation and *S. typhimurium* rdar morphotype and motility assay, residual DGC activity has been observed in different assays (Figure 2g and 4g) as it has been demonstrated that the ZMO1055 diguanylate cyclase activity activates cellulose biosynthesis (Li et al., 2023a). For example, the temporal appearance of a ‘brown’ morphotype might indicate an imbalance towards the diguanylate cyclase versus the phosphodiesterase activity by the activation of a novel matrix or cell membrane component alternatively indicate enhanced membrane permeability or other novel mechanisms that affect dye binding. Temporal alteration in the two opposite catalytic activities has been observed for other GGDEF-EAL domain proteins throughout the growth phase such as STM3388 in *S. typhimurium* (Kader et al., 2006). Thus, abolishment of the diguanylate cyclase activity of ZMO1055_ZM4_ would have been an alternative experimental set-up to assess the functionality of amino acid substitutions in the EAL domain.

Besides the direct functional impact of the A526V substitution, we observed that the ZMO1055 PAS-GGDEF-EAL protein in the phylogenetic context is subject to evolution. First of all, assessment of the 100 most homologous proteins indicated that the N-terminal sensory PAS domain shows the highest divergence (Figure S3b). It is however well known that sensory domains are usually subject to accelerated evolution. The GGDEF domain in homologous proteins is also subject to evolution although to a lesser degree. Closest *ZMO1055* homologs of member of the *Sphingomonas* genus are encoded at a conserved location on the chromosomes between *parC* encoding the DNA topoisomerase IV subunit A and *pcnB* coding for the poly(A)-RNA polymerase I in members of the Zymomonadaceae and the closely related Sphingomonadaceae family in contrast to other alpha-proteobacterial genera where the chromosomal context of ZMO1055 homologs is variable with homologous present as PAS-GGDEF-EAL or MHYT-PAS-GGDEF-EAL proteins (Figure S3c and d). Like in ZMO1055, the GGDEF motif in several of these homologues is subject to change including the presence of GADEF, AADEF and GGNEF variants, suggesting alterations in the degree of the catalytic activity, substrate specificity and/or loss of catalytic activity (Figure 3e). Of note, GGDEF domains with one amino acid substitution in the GGDEF motif has been shown to still possess catalytic activity (Cimdins-Ahne et al., 2023, Hunter, Severin et al., 2014, Perez-Mendoza, Coulthurst et al., 2011). Thus, an alteration in the GGDEF motif is not entirely predictive for a lack of catalytic activity.

While the EAL domain is predicted to be catalytically active in the vast majority of homologs (Figure S3d and e), the A526S, A526T, A526V and A526I substitution as present in ZMO1055 homologs can be considered as a first, still readily revertible, evolutionary steps towards a gradually reduced and subsequently abolished catalytic activity of the EAL domain. However, the impact of the A526 amino acid substitution might be specific to the ZMO1055 clade of EAL domains and/or requires a specific backbone constellation, as the sequence of distantly related EAL domain clades, such as for ZMO1487 is substantially different (Figure S3e). Eventually, ZMO1055 homologs can developed into proteins with catalytically incompetent GGDEF and EAL domains.

## Supporting information

Supplemental Information Figures and Experimental Procedures

## Acknowledgements

We thank Seyedmohammad Hosseinpourlamardi for the preparation of figure 6C. Lianying Cao received a one year visiting student scholarship from the Chinese Scholarship Council, CSC, to conduct research at the Karolinska Institutet, Stockholm, Sweden. Ute Römling received a grant from the Swedish Research Council for Natural Sciences and Engineering for cyclic di-GMP research. Lianyng Cao and Ute Römling are co-corresponding authors.

Data availability statement: All data are available within the manuscript.

The authors declare no conflict of interest.

