## Supplemental Information Figures and Experimental Procedures for "Previously uncharacterized aliphatic amino acid positions modulate the apparent catalytic activity of the EAL domain of ZMO_1055 and other cyclic di-GMP specific EAL phosphodiesterases"

*Corresponding authors

**Supplementary material**

**Title page	1**

**Index	2**

**Supplementary information	3**

**Supplementary Figure 1	5**

**Supplementary Figure 2	6**

**Supplementary Figure 3	7**

**Supplementary Table S1: Strains used in this study**

**Supplementary Table S2: Plasmids used in this study**

**Supplementary Table S3: Primers used in this study**

**Supplementary information**

**Materials and methods**

Protein sequences used for the alignment of GGDEF domain proteins in Figure 1a were: The sequences used in the alignment are PA0861_PSEAI (*Pseudomonas aeruginosa*); PA0575_PSEAI (*P. aeruginosa*); PleD_CAUCR, Q9A5I5 (*Caulobacter crescentus*); WspR_PSEAI, Q9HXT9 (*P. aeruginosa*); ARS29551.1 (A0A1X9YLB7; *Sphingomonas* sp. KC8); WP_084652956.1 (A0A1S1HAE2; *Sphingomonas haloaromaticamans*); TMJ20004.1 (A0A537MIH2; α-proteobacterium); ABQ66480.1 (A0A9J9H7P0; *Rhizorhabdus wittichii* DSM6014); TQL16391, (FBY51_1449, *Z. mobilis* NRRL B-4492); PROMIR_GGDEF (B3F0R5; *P. mirabilis*); STRGAL_GGDEF (WP_012961431.1; *S. gallolyticus* UCN34), YdeH_ECOLI, P31129 (*E. coli*); AdrA_SALTY, Q9L401 (*S. typhimurium*); PA4332_PSEAE, Q9HW69 (*P. aeruginosa*); DgcB_CAUCR, A0A0H3CAN8 (*C. crescentus*); CD1420_Cdif, Q18BU4 (*Clostridium difficile*); VCA0965_VIBCH, Q9KKY5 (*Vibrio cholerae*); ECA3270_PECAS, Q6D226 (*Pectobacterium atrosepticum*); GSU1658_GEOSL, Q74CL4 (*Geobacter sulfurreducens*); MXAN_2643, Q1D911 (*Myxococcus xanthus*); YciR_SALTY, A0A0F6B1Y8; DGC1_KOMXY, O87374 (*Komagataeibacter xylinus*); Y1354_MYCTU Rv1354c/P9WM13 (*Mycobacterium* *tuberculosis*); SE_0528_STAES, Q8CTF5 (*S. epidermidis*) and BifA_PSEAE, (Q9HW35; *P. aeruginosa*). In VCA0965, the sequence ‘RATNQHDY’ was deleted to optimize the alignment.

Protein sequences used for the alignment of EAL domain proteins in Figure 1b were: MorA_PSEAI (*P. aeruginosa*); PA0575_PSEAI (*P. aeruginosa*); RocR_PSEAI, Q9HX69 (*P. aeruginosa*); YciR_SALTY, A0A0F6B1Y8 (*S. typhimurium*); PDEA3_KOXYL, O87378 (*K. xylinus*); ARS29551.1 (A0A1X9YLB7; *Sphingomonas* sp. KC8); WP_084652956.1 (A0A1S1HAE2; *Sphingomonas haloaromaticamans*); TMJ20004.1 (A0A537MIH2; α-proteobacterium); ABQ66480.1 (A0A9J9H7P0; *Rhizorhabdus wittichii* DSM6014); TQL16391, (FBY51_1449, *Z. mobilis* NRRL B-4492); YahA_ECOLI, P21514 (*E. coli* K-12); YhjH_SALTY, A0A0F6B886 (*S. typhimurium*); YE2225_YENTE, A1JQ37 (*Yersinia enterocolitica*); DGC2_Komxyl, O87377 (*K. xylinus*); LapD_PSEFL, Q3KK31 (*P. fluorescens* Pf0-1); FimX_PSEAI, Q9HUK6 (PA4959; *P. aeruginosa*); STM1344_SALTY, D0ZW85 (*S. typhimurium* ATCC14028); ToxR_PSEAI, P09852 (*P. aeruginosa*); CsrD_ECOLI, P13518 (*E. coli* K-12).

Protein sequences used in the phylogenetic tree of Figure S3e were: MBP2137887.1, *Sphingomonas* *echinoides* BE319; WP 010215794.1, *Sphingomonas* sp. PAMC 26621; SDA14076.1, *Sphingomonas* sp. NFR15; MBN8814462.1; *Sphingomonas* sp. SCN18_26_2_15_R5_F_65_27; OQW73042.1, Proteobacteria bacterium ST_bin13; MBA4773209.1, *Sphingomonas* sp. MCMED-G21; TCP36799.1, *Sphingomonas* sp. BK235 EV292_101297; OWK33834.1, *Sphingomonas dokdonensis*; KKI18272.1, *Sphingomonas* sp. Ag1; PKP94256.1, HGW-Alphaproteobacteria-16; KRC82426.1; *Sphingomonas* sp. Root241; PVX62630.1, *Sphingomonas* sp. CF311; WP 197418082.1, *Sphingomonas* sp. CCH13-B11; PZO75509.1, *Sphingomonas hengshuiensis*; KQM18664.1, *Sphingomonas* sp.; WP 171744425.1, *Sphingomonas* sp. AP4-R1; WP 167072910.1, *Sphingomonas vulcanisoli*; TMJ20004 1, Alphaproteobacteria bacterium AP_24; MBE1481263.1; *Sphingomonas* sp. OAS965; ARS29551 1 SPH KC8, *Sphingomonas* sp. KC8; WP 084652956 1 SPHHAL; *Sphingomonas*; ABQ66480 1 SPHWIT; *Sphingomonas wittichii* RW1; WP_013933534.1 (F8EU54, *Zymomonas mobilis* subsp. pomaceae ATCC29192) and WP 163642225.1, *Methylobacterium* sp. BTF04; WP 056485266.1, *Methylobacterium* sp. Leaf117; TNC10695.1. *Methylobacterium* sp. 17Sr1-39; WP 060851252.1, *Methylobacterium aquaticum*; WP 090962595.1, *Aureimonas phyllosphaerae*; SPZ47856_PSEAI.1, *Agrobacterium tumefaciens* biovar 1; RYD45316.1, Sphingomonadales bacterium; MBO6856075.1, *Roseibium* sp. JJ875_04115; GBE42618.1, bacterium *BMS3Bbin10;* WP_154738706.1, *Hyphomicrobium* sp. Xq; MBL8709276.1, *Rhodospirillaceae* bacterium; WP 133292192.1, *Dankookia rubra* PRK10060 and STM3615, *Salmonella typhimurium* ATCC14028; STM0468, S. *typhimurium* ATCC14028; PA3258, *P. aeruginosa* PAO1.

**Figures**

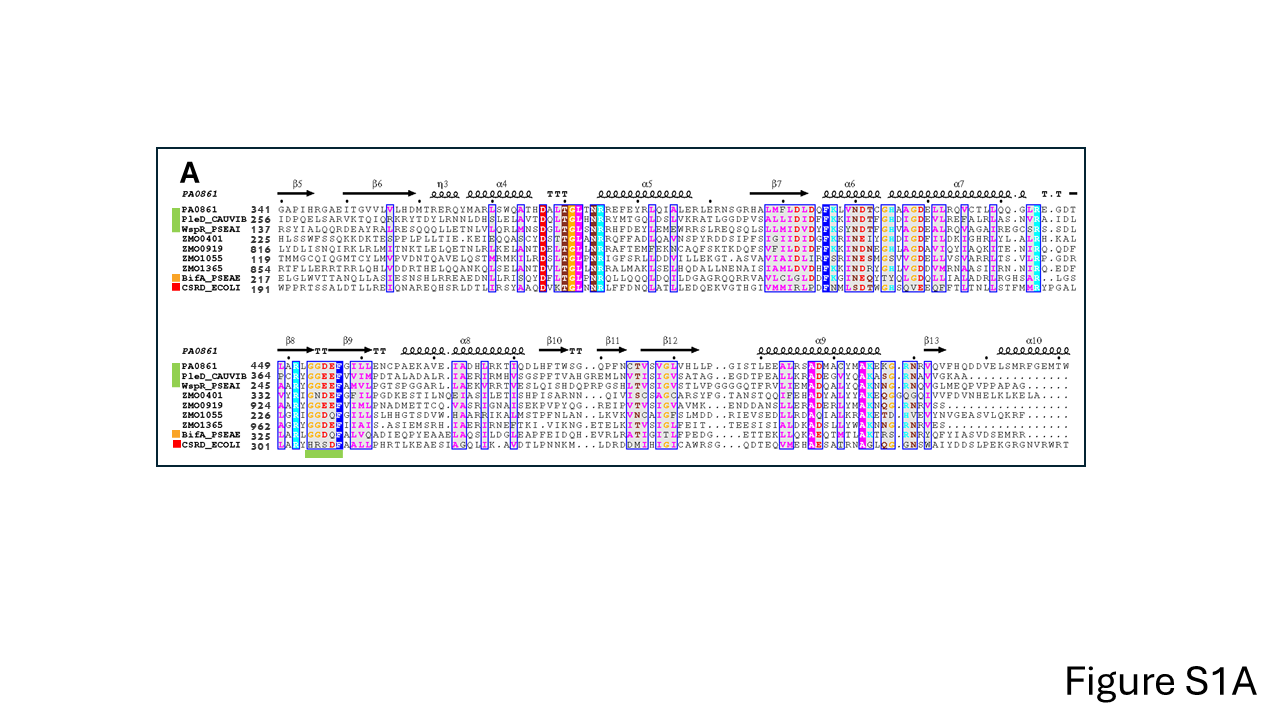

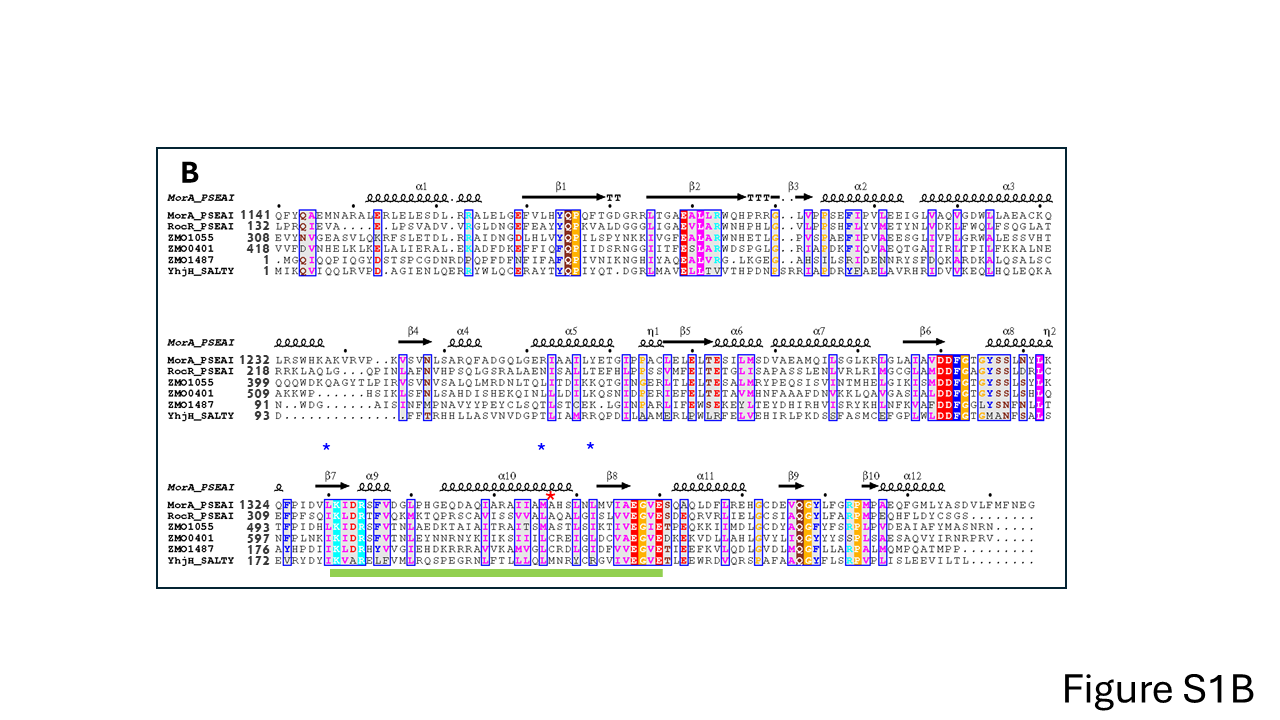

Figure S1: Alignment of the GGDEF and EAL domains encoded by the *Zymomonas mobilis* ZM4 genome with catalytically functional reference GGDEF and EAL domain proteins.

Figure S1a: Alignment of all GGDEF domains of *Z. mobilis* ZM4 with template and reference GGDEF domains. The GGDEF domain of the diguanylate cyclase PA0861 of *P. aeruginosa* (PDB: 5XGD) has been used for secondary structure designation. Green side bar indicates catalytically active domains (PleD and WspR), yellow side bar catalytically functional with C-terminal EAL domain (BifA) and red side bar catalytically inactive domaisn (CsrD). Underlined by a green bar is the GG(D/E)EF catalytic motif.

Figure S1b: Alignment of all EAL domains of *Zymomonas mobilis* with reference EAL domains. The EAL domain of the phosphodiesterase MorA of *P. aeruginosa* (PDB: 4RNI) has been used for secondary structure designation. RocR and the phosphodiesterase YhjH were added as comparison representing class I and class II phosphodiesterases (see Figure 1 a and b for explanation). All reference EAL domains are active phosphodiesterases. Underlined in green is the amino acid sequence between the conserved K(I/L/V)D and the EG(V/I)E motif that contains A526 (indicated by a green star). Blue stars indicate amino acids 499, 525 and 531 substituted in the course of this work.

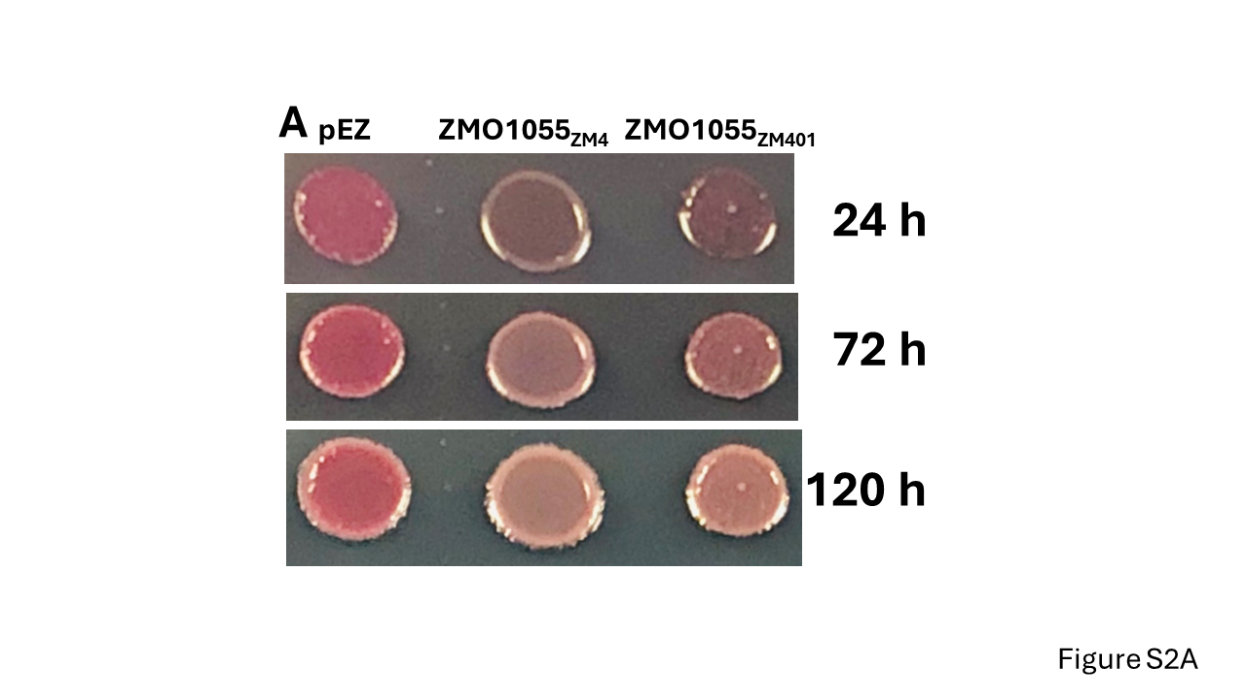

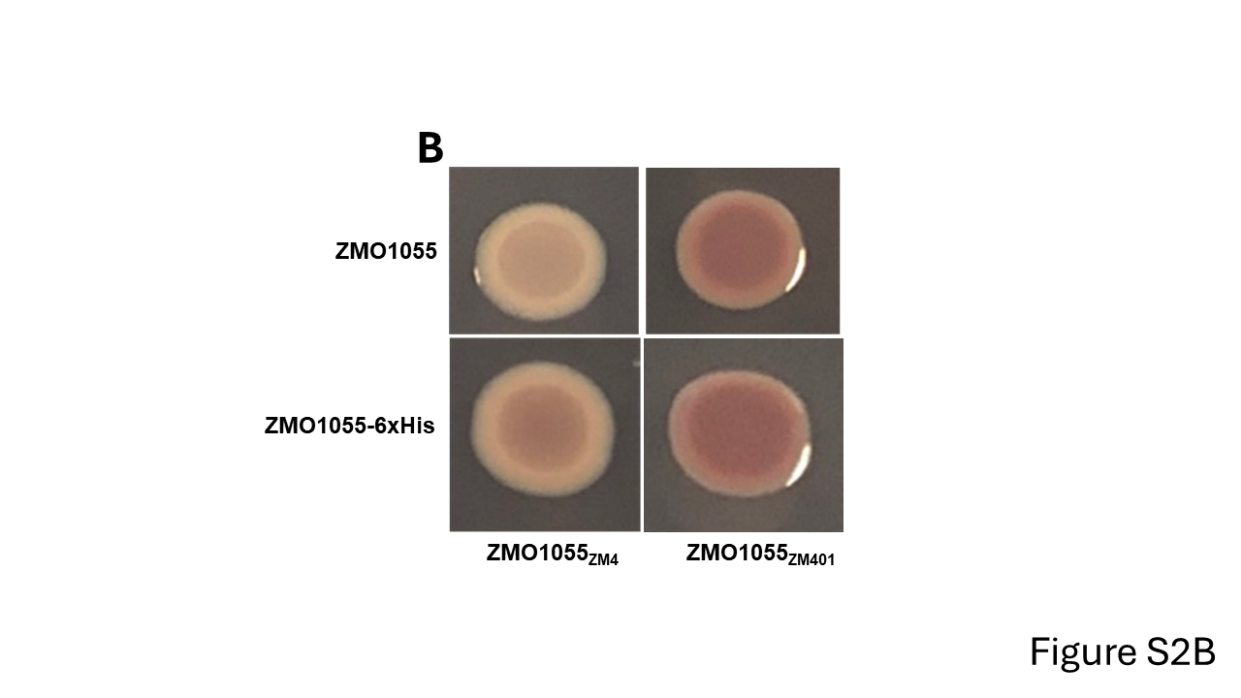

Figure S2: Colony morphologies of *Z. mobilis* ZM401 upon prolonged incubation up to 72 h (a) and effect of 6xHis C-terminal of ZMO1055 on colony morphology of *S. typhimurium* UMR1 Δ*yhjH* (b). (a) *Z. mobilis* ZM401 with pEZ vector control, ZMO1055_ZM4_ and ZMO1055_ZM401_ cloned in pEZ. Cells were grown on Congo red RM agar plates at 28°C for indicated time points. (b) *S. typhimurium* UMR1 Δ*yhjH* with ZMO1055_ZM4_ and ZMO1055_ZM401_ cloned in pEZ without and with a C-terminal 6xHis-tag. Cells were incubated on Congo red LB without salt agar plates at 28°C for 24 h.

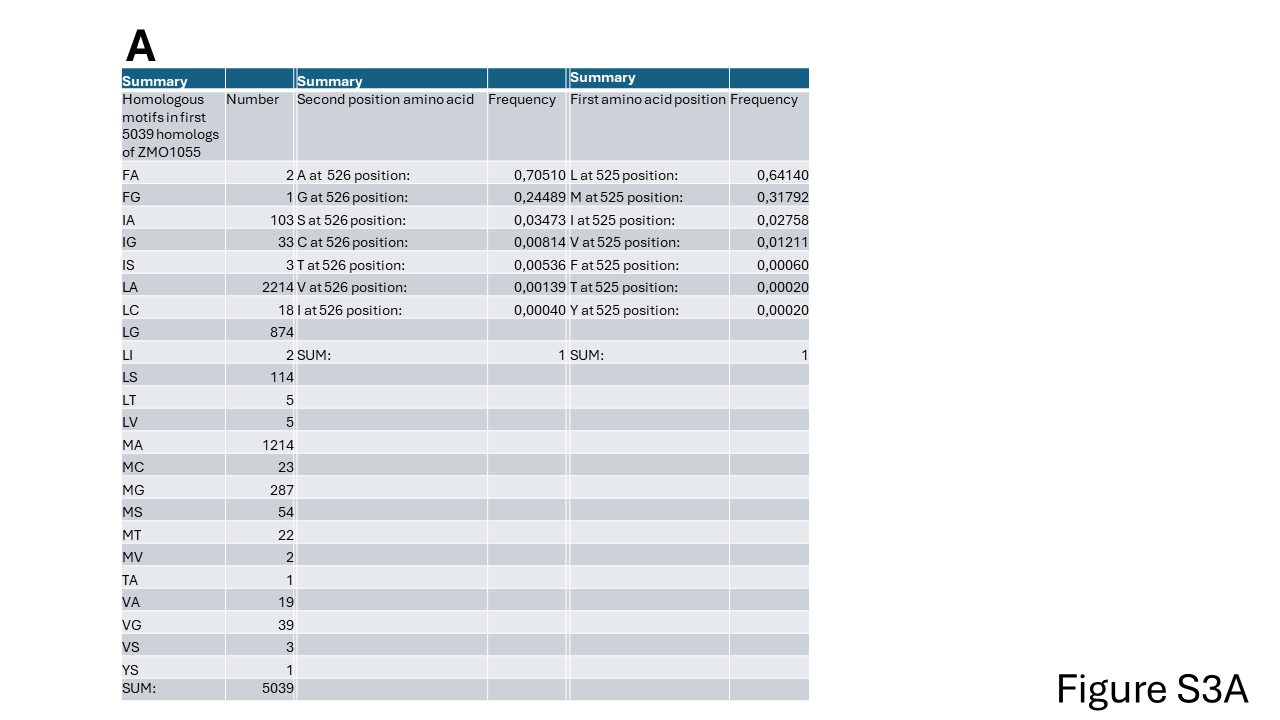

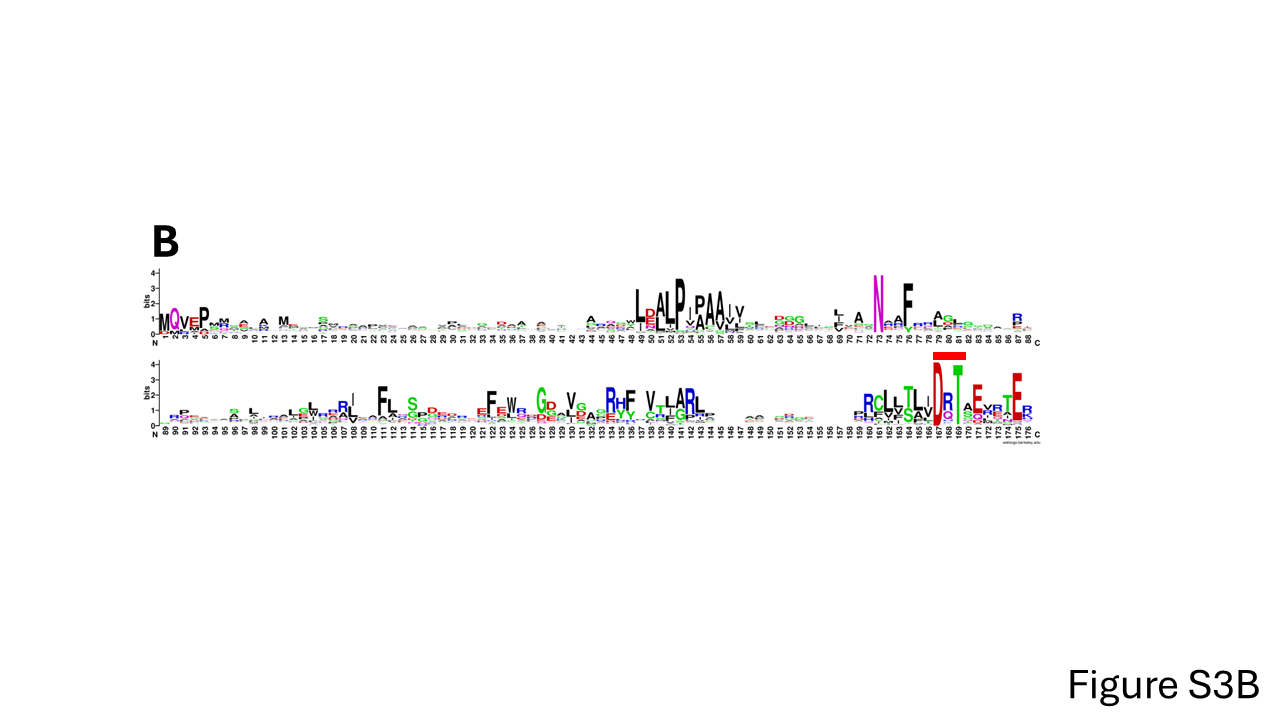

Figure S3: Bioinformatic analysis of the PAS-GGDEF-EAL protein ZMO1055_ZM4_ in the context of the most similar homologs.

Figure S3a: Frequency of amino acids at the M_525_A_526_ position. The 5039 most similar non-redundant ZMO1055_ZM4_ homologs were retrieved by Blast (April 2021), aligned with subsequent manual curation and the frequency of distinct amino acids at the M_525_A_526_ position calculated.

Figure S3b: Sequence logo of the ZMO1055_ZM4_ clade PAS domain. The PAS domain of the 100 most similar PAS-GGDEF-EAL domain proteins as retrieved by Blast (April 2021) has been aligned and a WebLogo has been created. Indicated by a red bar is the C-terminal end of the PAS domain (DxT motif).

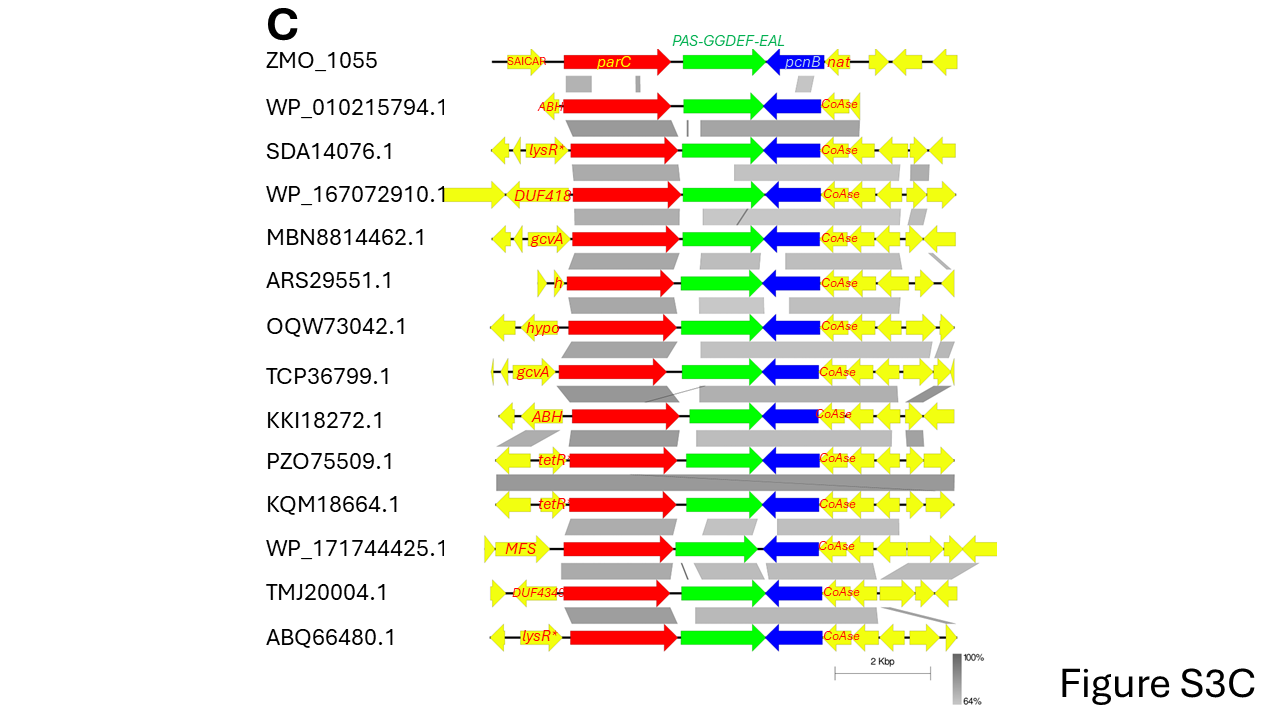

Figure S3c: Chromosomal position of the ZMO1055_ZM4_ homologs from the genus *Sphingomonas*. The immediate chromosomal position of most similar homologs of ZMO1055_ZM4_ from genus *Sphingomonas* representatives (with reference to the phylogenetic tree in Figure S3e).

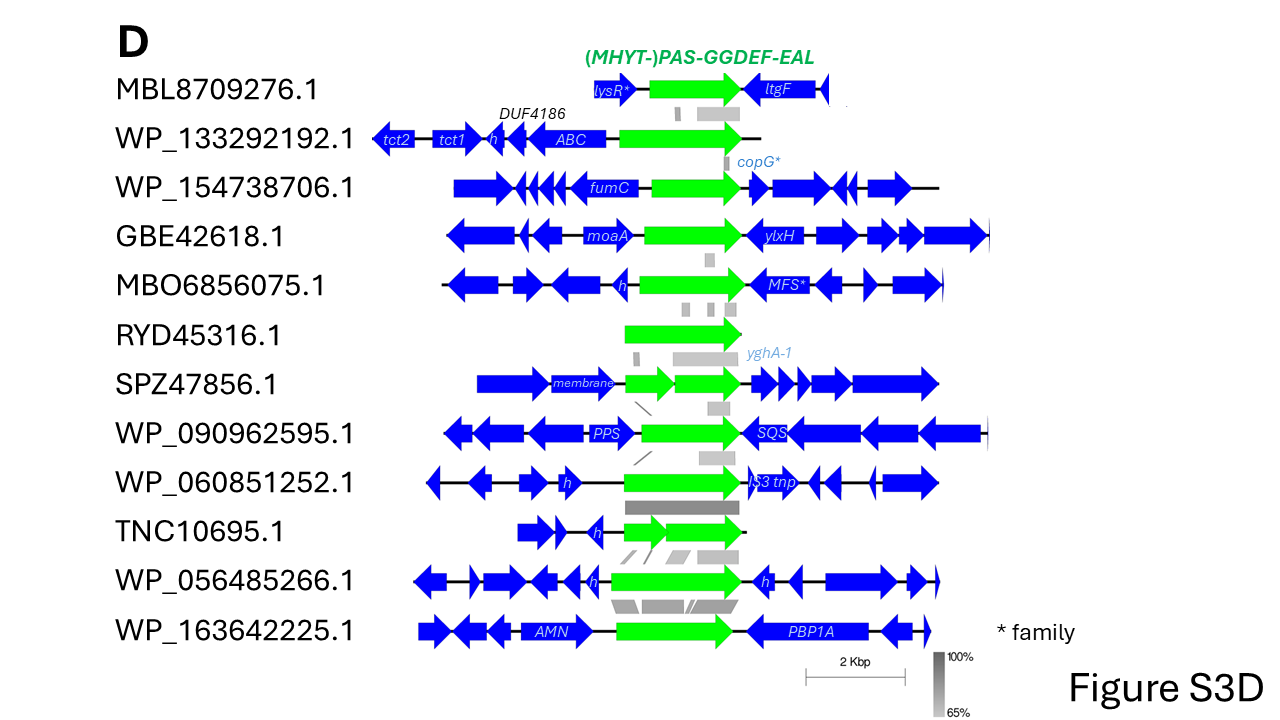

Figure S3d: Chromosomal position of the ZMO1055_ZM4_ homologs outside the genus *Sphingomonas*. The immediate chromosomal position of most similar homologs of ZMO1055_ZM4_ outside the genus *Sphingomonas* (with reference to the phylogenetic tree in Figure S3e)..

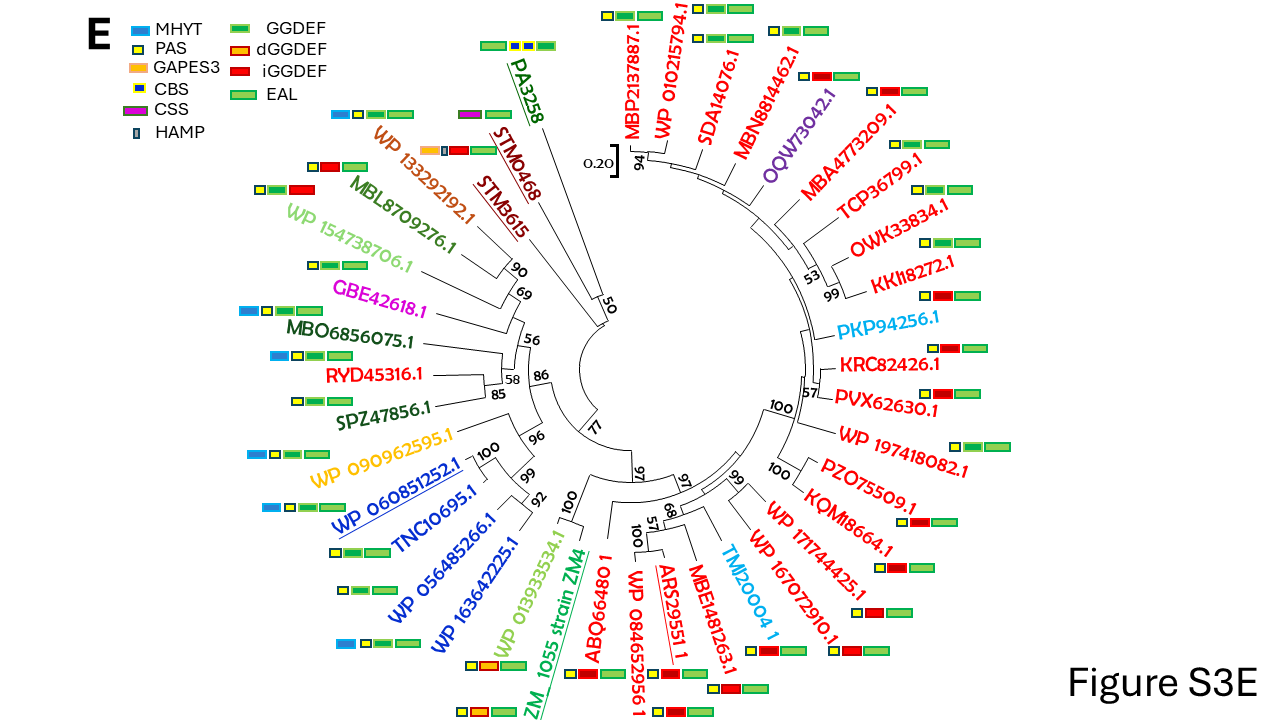

Figure S3e: Phylogenetic tree of the EAL domain from most similar homologous GGDEF-EAL proteins to the ZMO1055_ZM4_ EAL domain as retrieved by Blast (April 2021) from the NCBI database and the proteins used to create the A526V equivalent. The EAL domains of representatives of most similar PAS-GGDEF-EAL domain proteins have been aligned with subsequent manual curation and a Maximum-Likelyhood phylogenetic tree with 1000 bootstraps has been constructed in MEGA 7.0. Next to the protein designation, the domain structure of the respective protein is given. Proteins used for mutational analysis are underlined. Symbols in the upper left corner indicate the different signaling and catalytic domains. GGDEF and EAL are predicted catalytically active diguanylate cyclase and phosphodiesterase domains. dGGDEF indicates degenerated GGDEF motif, but catalytically active domain as experimentally demonstrated. iGGDEF indicates degenerated GGDEF or other signature motif with no experimental documentation of catalytic activity. MHYT, PAS (Per-ARNT-Sim), GRAPE3, CBS (cystathionine-beta-synthase), CSS and HAMP (present in Histidine kinases, Adenyl cyclases, Methyl-accepting proteins and Phosphatases) are N-terminal signaling domains.

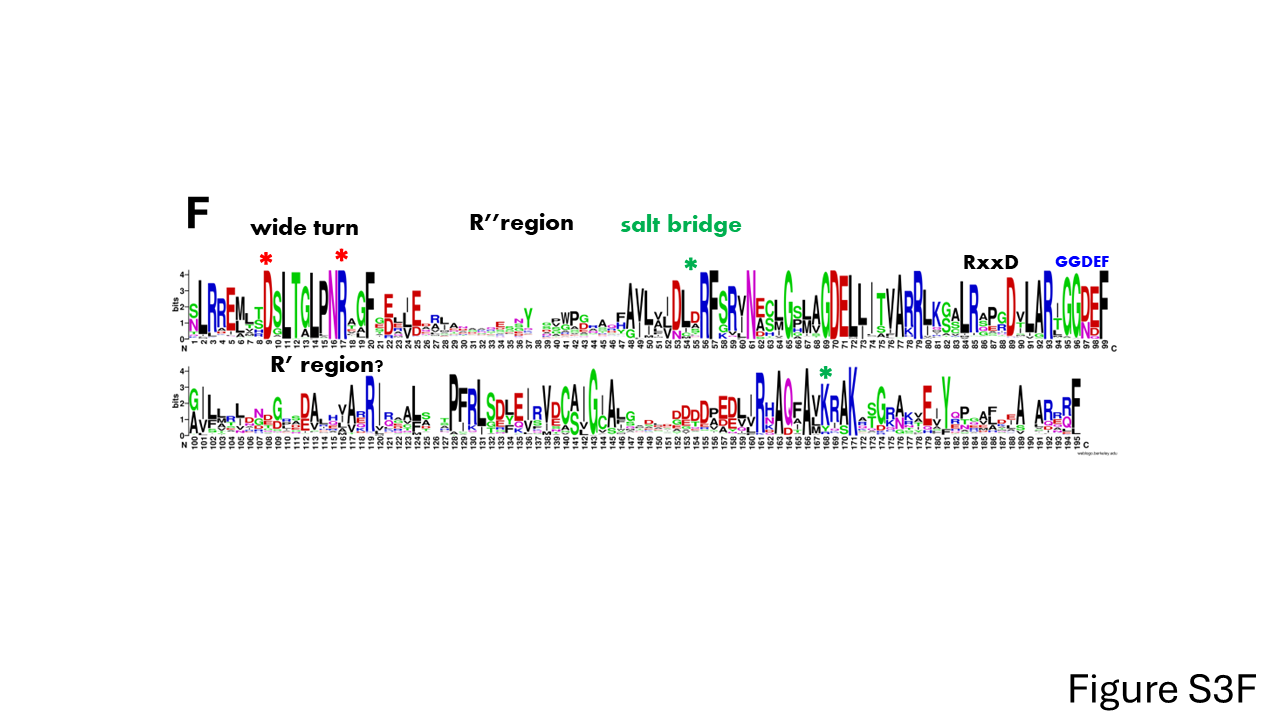

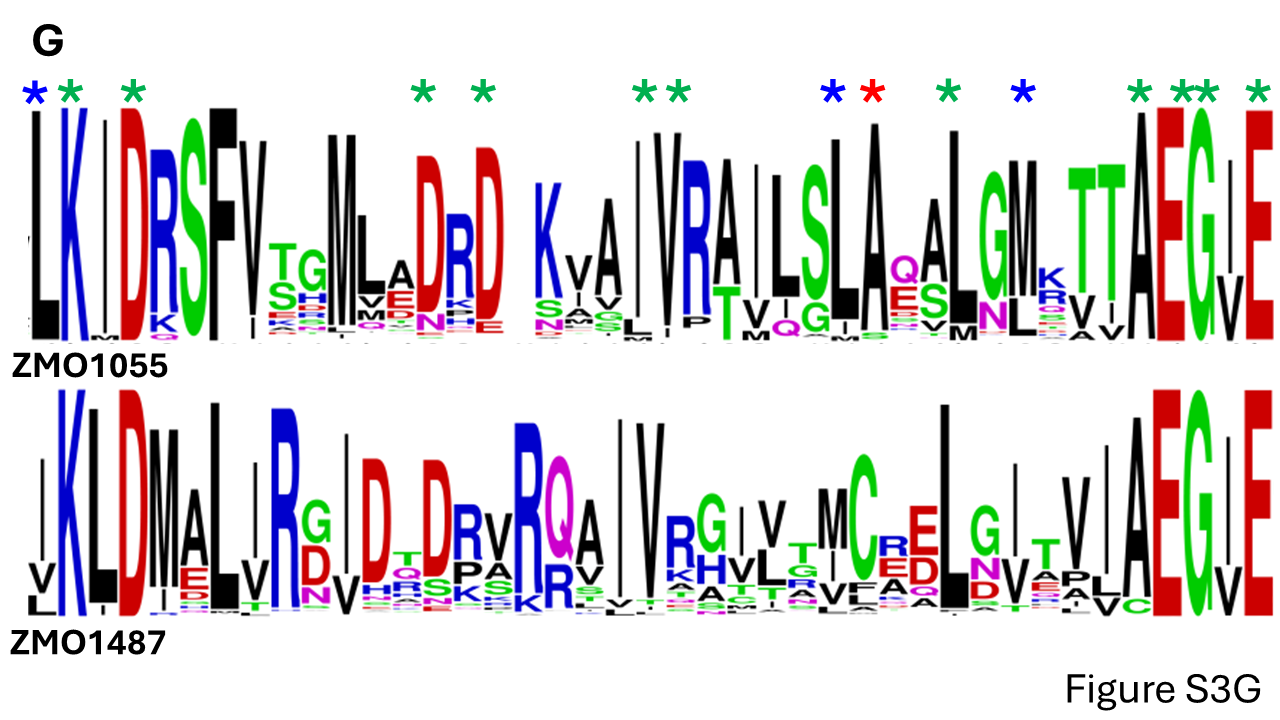

Figure S3f: Sequence logo (WebLogo) of the ZMO1055_ZM4_ clade GGDEF domains. The GGDEF domains from the most similar homologous GGDEF-EAL proteins to ZMO1055_ZM4_ as retrieved by Blast (April 2021) from the NCBI database. The GGDEF domain of the 1000 most similar (MHYT)-PAS-GGDEF-EAL domain proteins as retrieved by Blast has been aligned with subsequent manual curation and a sequence logo has been constructed. Functionally relevant motifs such as the catalytic site GGDEF motif, the allosteric I-site RxsD motif, the wide turn, the R’ and R’’ region, the wide turn and the salt bridge are indicated.

Figure S3g: Sequence logo of the amino acid sequence between the conserved KID and EGxE motif of ZMO1055_ZM4_ and ZMO1487_ZM4_ clade EAL domains. One thousand EAL domains of the most similar homologous GGDEF-EAL proteins to the ZMO1055_ZM4_ GGDEF domain and 1000 EAL domains of the most similar homologous proteins to ZMO1487_ZM4_ have been retrieved by Blast (October 2021) from the NCBI database. The EAL domains were aligned with subsequent manual curation and a sequence logo has been retrieved displaying the amino acid sequence conservation between the conserved KID and EGxE motifs. Red star, position A526 in ZMO1055; blue stars, amino acids mutated in this study; green stars, reference amino acids highly conserved between ZMO1055 and ZMO1487 closest homologues.

**References**
